# Impaired bone morphogenetic protein signaling pathways disrupt decidualization in endometriosis

**DOI:** 10.1101/2023.09.21.558268

**Authors:** Zian Liao, Suni Tang, Peixin Jiang, Ting Geng, Dominique I. Cope, Timothy N. Dunn, Joie Guner, Linda Alpuing Radilla, Xiaoming Guan, Diana Monsivais

## Abstract

It is hypothesized that impaired endometrial decidualization contributes to decreased fertility in individuals with endometriosis. To identify the molecular defects that underpin defective decidualization in endometriosis, we subjected endometrial stromal cells from individuals with or without endometriosis to time course *in vitro* decidualization with estradiol, progesterone, and 8-bromo-cyclic-AMP (EPC) for 2, 4, 6, or 8 days. Transcriptomic profiling identified differences in key pathways between the two groups, including defective bone morphogenetic protein (BMP)/SMAD4 signaling (*ID2, ID3, FST*), oxidate stress response (*NFE2L2, ALOX15, SLC40A1*), and retinoic acid signaling pathways (*RARRES, RARB, ALDH1B1*). Genome-wide binding analyses identified an altered genomic distribution of SMAD4 and H3K27Ac in the decidualized stromal cells from individuals without endometriosis relative to those with endometriosis, with target genes enriched in pathways related to signaling by transforming growth factor β (TGFβ), neurotrophic tyrosine kinase receptors (NTRK), and nerve growth factor (NGF)-stimulated transcription. We found that direct SMAD1/5/4 target genes control FOXO, PI3K/AKT, and progesterone-mediated signaling in decidualizing cells and that BMP2 supplementation in endometriosis patient-derived assembloids elevated the expression of decidualization markers. In summary, transcriptomic and genome-wide binding analyses of patient-derived endometrial cells and assembloids identified that a functional BMP/SMAD1/5/4 signaling program is crucial for engaging decidualization.

## INTRODUCTION

Endometriosis is a debilitating disease affecting ∼ 190 million women of reproductive age globally^1^. Defined as the occurrence of endometrial glands and stroma compartments outside of the uterine cavity, endometriosis leads to inflammatory conditions of the lesion sites. Lesions are typically located within the pelvic areas but can also involve the bowel, bladder, diaphragm, pleura, and lungs, and other distant sites^2^. Patients often suffer from chronic pelvic pain, severe dysmenorrhea, or infertility, which can disrupt the social and professional life of affected patients. Currently, there is no definite explanation for the pathogenesis of endometriosis; however, several theories have been proposed to explain this multifactorial process. In 1925, Sampson introduced the idea of retrograde menstruation, where endometrial debris can travel backward through the Fallopian tubes and into the pelvic and peritoneal cavities during menses^3^. However, studies have shown that around 90% of women experience retrograde menstruation and the majority do not develop into endometriosis^4,5^. Such observations give rise to alternative theories that posit the origin of the disease is from non-uterine tissues. Sources of transformed ectopic cells include bone marrow (mesenchymal and hematopoietic stem cells)^6^, müllerian duct remnants^7^ and coelomic metaplasia^8^. Regardless of their initial pathogenesis, the main symptomatic process involves increased production of inflammatory cytokines and pain mediators, as well as dysfunction of sympathetic nerve fibers^9–11^. Treatment options for endometriosis are limited to empirical nonsteroidal anti-inflammatory drugs (NSAIDs), hormonal therapies or surgery^5^. Moreover, evidence has shown that surgical diagnosis of endometriosis was correlated with an increased risk of ovarian cancer^12^.

In addition to causing pain and inflammation, endometriosis often leads to infertility^13–15^. Around 40 percent of women with endometriosis are estimated to have infertility^16,17^, and of women with infertility, 25 to 50 percent are estimated to also suffer from endometriosis^18,19^. Endometriosis affects fecundity mainly by impairing ovarian functions, inducing chronic intraperitoneal inflammation and progesterone resistance. Patients with endometriosis present with an abnormally prolonged follicular phase^20^, which further leads to dysfunctional folliculogenesis and granulosa cell cycle kinetics^21,22^. As a key feature of endometriosis, chronic intraperitoneal inflammation stems from increased levels of inflammatory cytokines, chemokines as well as prostaglandins. Such inflammatory processes can lead to infertility by decreasing intrafollicular estrogen level^23^, oocytes quality^24^ and sperm motility^24^.

The normal function of the eutopic endometrium is also compromised in patients with endometriosis, as demonstrated by progesterone resistance accompanied by declined expression of progesterone receptor (PR) and coactivators^25,26^. Deficient progesterone signaling pathways likely lead to impaired decidualization, defective embryo implantation and increased infertility rates in patients with endometriosis^27^. Additional signaling defects have been identified in the endometrium of women with endometriosis, including defective mesenchymal stem cell differentiation^28,29^, increased decidual senescence and elevated pro-inflammatory stress^30^. Uncovering the mechanisms and pathways involved that cause the negative effect of endometriosis on the eutopic endometrium is critical to help optimize chances for successful pregnancy.

Our studies aim to uncover the molecular underpinning of infertility associated with endometriosis by focusing on the effects of the human endometrium. Here, we use patient-derived stromal cells and state-of-the-art endometrial stromal and epithelial assembloids to define key signaling alterations during decidualization in patients with endometriosis.

## RESULTS

### Transcriptomic profiling in endometrial stromal cells from individuals with and without endometriosis reveals activation of key pathways during *in vitro* decidualization

The human endometrium undergoes spontaneous decidualization in response to the rising levels of the ovarian hormone, progesterone^31^. The concerted action of estrogen and progesterone transforms the endometrium from a non-receptive state into a receptive state, subsequently allowing embryo implantation and development to occur. Because patients with endometriosis can have worsened fecundity due to defective endometrial function, we aimed to systematically determine the transcriptomic differences between the two groups during *in vitro* stromal cell decidualization.

We used a well-characterized method,^32–35^ to induce endometrial stromal cell decidualization and to compare differentially expressed genes between normal and endometriosis samples at different stages after the EPC treatment, from early (2 days) to late (8 days) decidualizing stages. Patient-derived endometrial stromal cells from patients with and without endometriosis were cultured *in vitro* and treated with estrogen (E2), medroxyprogesterone acetate (MPA) and 8-bromo cyclic adenosine monophosphate (cAMP) (EPC) to induce decidualization *in vitro*, and then collected at 2, 4, 6, 8 days after EPC cocktail treatment to profile transcriptomic changes using RNA sequencing (Figure 1A). We performed a time course comparison between the differentially expressed genes at each timepoint respectively. In total, 334 transcripts changed significantly during the EPC treatment in the cells derived from individuals without endometriosis (n=3, “normal”), with 152 transcripts showing an increase and 182 showing a decrease by day 8 of EPC treatment (Supplementary Table 1, >1.4, <0.4-fold change, FDR < 0.05). In stromal cells from individuals with endometriosis (n=4), 878 transcripts showed a significant change after EPC treatment, with 464 being increased and 414 decreased by Day 8 of EPC treatment (Supplementary Table 1, >1.4, <0.4-fold change, FDR < 0.05). Among these, only 122 transcripts (12.4%) were shared between normal and endometriosis upregulated genes, and 105 (10.7%) shared genes were conserved in the downregulated genes between the two groups (Supplementary Figure 1A). These results indicate that endometrial stromal cells arising from individuals with endometriosis display a different transcriptomic response to decidualization treatment *in vitro*.

**Figure 1.**
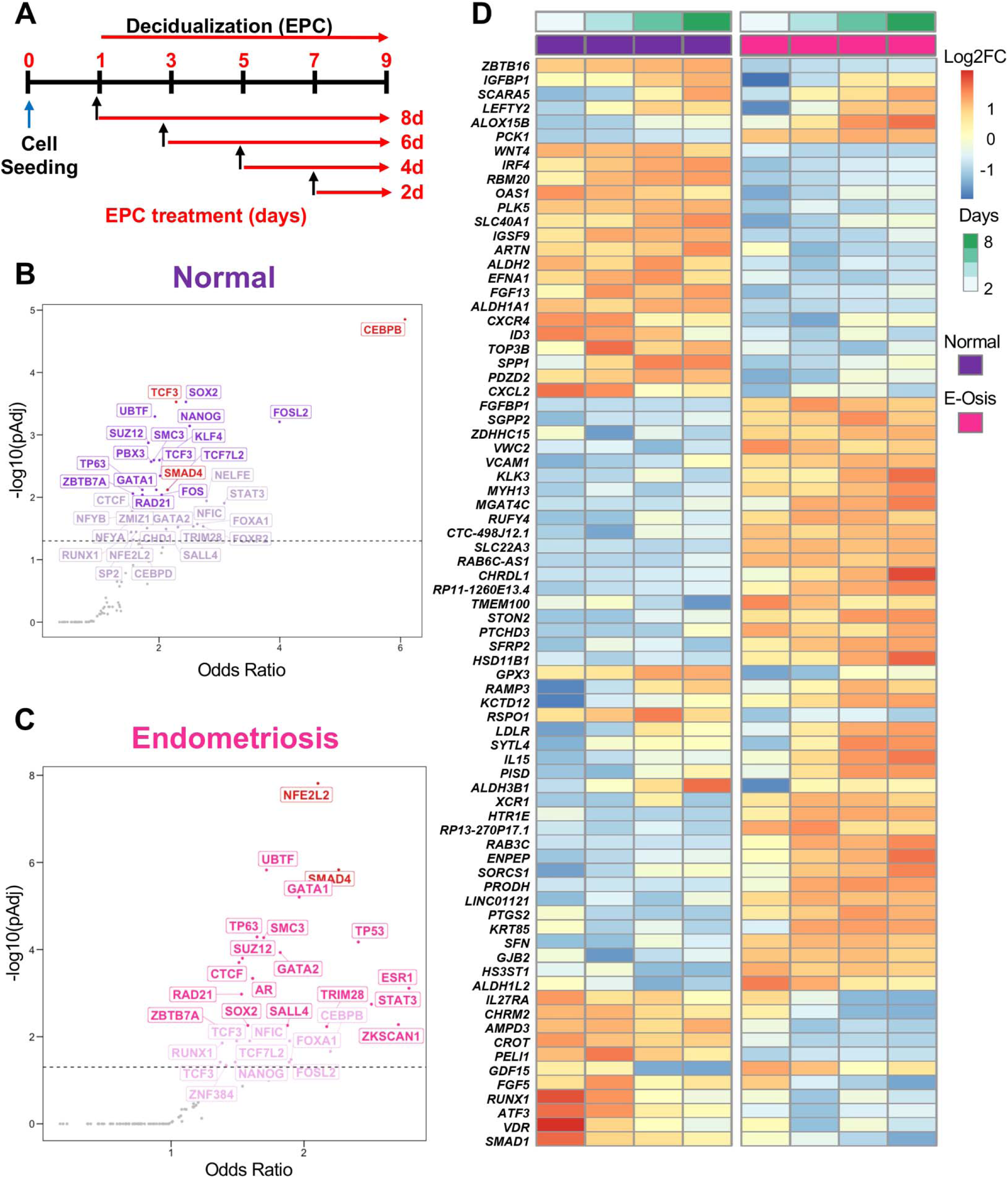
Transcriptomic profiling of endometrial stromal cells from individuals with or without endometriosis reveals key differences during *in vitro* decidualization. A) Primary endometrial stromal cell cultures from individuals without (n=3, “normal”) or with endometriosis (n=4) were subjected to a time-course decidualization treatment. After plating, cells were treated with vehicle or with the decidualization cocktail (35nM estradiol, 1µM medroxyprogesterone acetate, 50µM cAMP, “EPC”) for 2, 4, 6, or 8 days. RNA sequencing was performed and the decidualization response within normal and endometriosis stromal cells was determined by normalizing differentially expressed genes relative to the Day 0 (vehicle)-treated cells. B-C) Upstream transcriptional regulators were identified by searching for conserved ENCODE and ChEA consensus gene targets among the differentially expressed genes in the normal (B) and endometriosis (C) groups. CEBP/β and TCF3 emerged as top transcription regulators for normal decidualizing cells (B), while NFE2L2 and SMAD4 were determined to be major upstream regulators for endometriosis. D) Heatmap displays gene expression over time within the normal and endometriosis (“E-Osis”) groups treated with EPC using normalized z-scores. Color represents log2 fold-change relative to baseline (day 0).

To understand the pathways that were overrepresented among the differentially expressed genes, we performed a gene ontology analysis of all the differentially regulated genes (>1.4, <0.4-fold change, FDR < 0.05) in the normal or endometriosis datasets (Supplementary Table 2). Among the top 10 categories in the normal patients, we found that cytosolic tRNA aminoacylation, Interleukin-2 signaling, TGFβ-regulation of extracellular matrix and BDNF signaling pathways were overrepresented among the differentially expressed genes at day 8 of EPC relative to baseline (Day 0) (Supplementary Figure 1B). Enriched pathways in the endometriosis dataset included categories such as, TGFβ-regulation of extracellular matrix, interleukin-2 signaling, cardiomyocyte hypertrophy, FSH regulation of apoptosis, and Leptin signaling pathway (Supplementary Figure 1C). Our results suggest that while the cells from both groups displayed uniquely regulated genes, several of these key pathways, such as TGFβ signaling and interleukin-1 signaling were shared between the two groups.

We performed an upstream transcription factor analysis of the differentially regulated genes in the endometriosis or normal stromal cells to identify master regulatory networks driving the differential transcriptional response (Supplementary Table 3). By mining the consensus gene targets in the ENCODE and ChEA Transcription Factor Targets dataset^36,37^, we identified that genes regulated by the CCAAT Enhancer Binding Protein Beta (CEBP/β) and Transcription Factor 3 (TCF3) were enriched in the normal stromal cells (Figure 1B). CEBP/β has been shown to be a key factor in endometrial stromal cell decidualization that controls the transcription of the PR^38,39^ TCF3 is also shown to control endometrial stromal cell proliferation and decidualization^40^. On the other hand, regulation of genes by the NFE2 Like BZIP Transcription Factor 2 (NFE2L2) and SMAD4 transcription factors was highly enriched in the endometriosis dataset (Figure 1C). NFE2L2 is an important regulator of oxidative stress response that controls the expression of genes that contribute to ferroptosis resistance^41–43^. SMAD4 is the downstream activated transcription factor controlling expression of genes downstream of bone morphogenetic proteins (BMPs, through SMAD1 and SMAD5) or TGFβ/activin ligands (through SMAD2 and SMAD3)^44^. Some of these differential responses could also be observed at the gene level (Figure 1D), as indicated by the expression of *IGFBP1*^45^*, ZBTB16*^46^ (decidualization markers), *SLC40A1, GPX3, PTGS2*^47,48^ (markers of oxidative stress, ferroptosis markers), or *LEFTY2, SMAD1*^49^ (BMP/SMAD-signaling pathways). In summary, while conventional transcriptional programs driving decidualization were found to be overrepresented in the endometrial stromal cells from individuals without endometriosis, endometrial stromal cells from individuals with endometriosis showed a transcriptional program suggesting a perturbation in oxidative stress and BMP/TGFβ signaling pathways.

### Identification of perturbed BMP/TGFβ signaling pathways in the decidualizing stromal cells from individuals with endometriosis

We examined the dynamic profiles in the early and late decidual cell transcriptomes and visualized the differentially expressed genes between individuals with and without endometriosis in volcano plots using a >2 or < ½ fold-change and FDR < 0.05 (Figure 2 A-D). At baseline, we observed that 393 transcripts were differentially expressed between the normal and endometriosis groups, with 166 being upregulated and 227 downregulated in the endometriosis group compared to the normal group. Two days after EPC administration, 400 genes were down regulated in the endometriosis samples compared to the normal counterparts. We then observed 216, 95 and 99 genes to be down regulated on Day 4, Day 6, and Day 8, in endometriosis samples compared to the normal samples, respectively. Meanwhile, 323, 138, 64 and 43 genes were up regulated in endometriosis compared to normal at Day 2, Day 4, Day 6, and Day 8, respectively (Supplementary Table 4).

**Figure 2.**
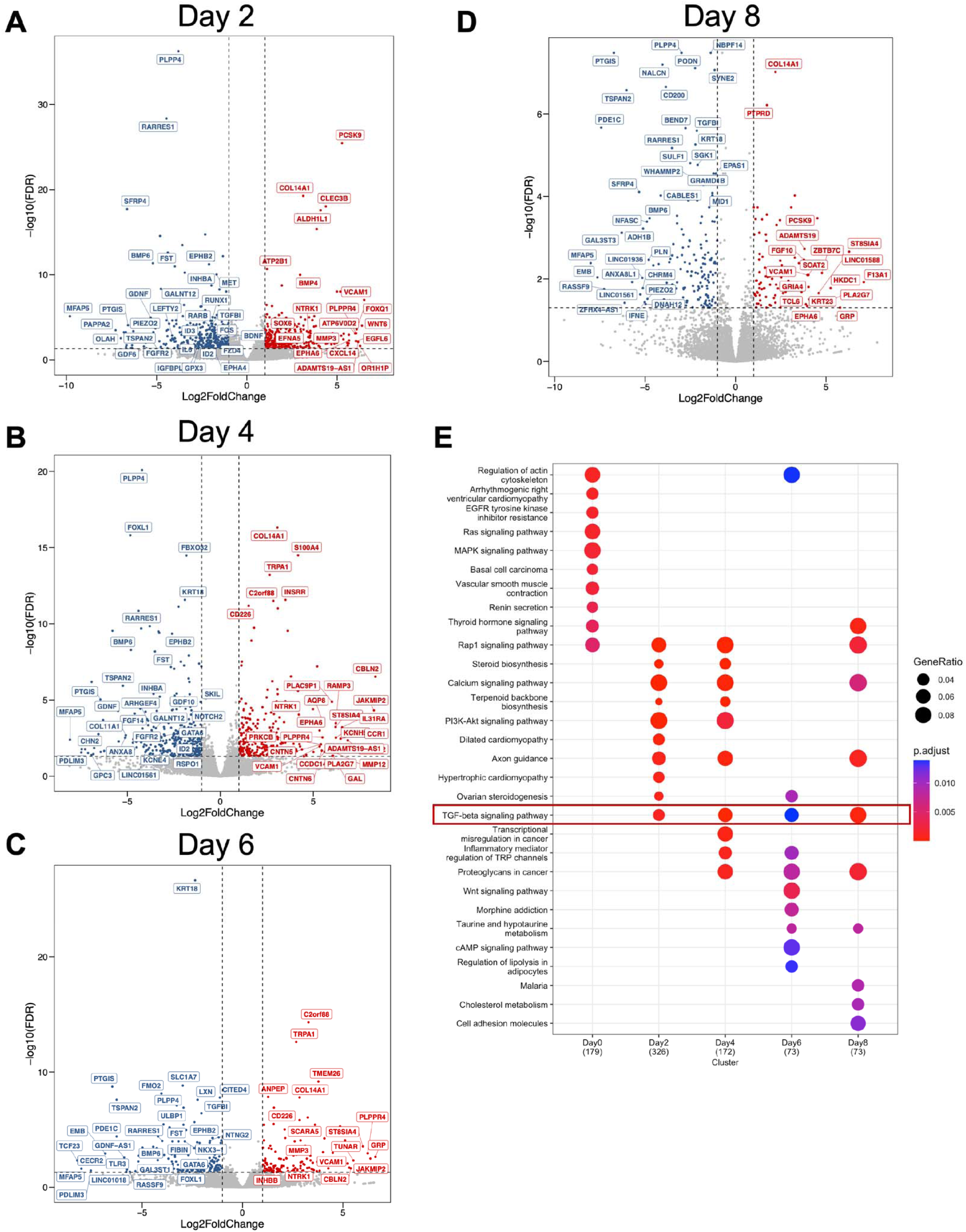
BMP/TGFβ signaling pathways are defective in the decidualizing stromal cells from individuals with endometriosis. Differentially expressed genes between the endometrial stromal cells of individuals without (n=4) or with endometriosis (n=3) at each time point during the decidualization treatment were identified and visualized as volcano plots. Differentially expressed genes were determined using a cut off (|log2 fold-change| >1 and FDR < 0.05, red denotes increased genes, blue denotes decreased genes, gray indicates no significant change) and displayed following Day 2 (A), Day 4 (B), Day 6 (C), or Day 8 (D) of treatment with the EPC decidualization cocktail (35nM estradiol, 1µM medroxyprogesterone acetate, 50µM cAMP). E) Gene ontology analysis of the differentially expressed genes was performed at each time point and visualized as a Dot Plot. Genes in the TGFβ signaling pathway were identified to be enriched at each of the time points after EPC treatment.

To understand the pathway dependent differences in the endometriosis groups, we implemented Kyoto Encyclopedia of Genes and Genomes (KEGG) pathway enrichment analysis in the differentially expressed genes spanning from Day 0 to Day 8 timepoints between normal and endometriosis donors (Figure 2E). Our enrichment analysis showed distinct enrichment patterns at each of the different time points we assessed. Of notice, the TGFβ signaling pathway was the only category shared in the timepoints during decidualization. After overlapping the differentially expressed genes from different time points, we identified that 48 genes were consistently down-regulated and that 20 genes were consistently up-regulated regardless of the EPC treatment length (Supplementary Figure 2A-B). Further enrichment for genes associated with human diseases by DisGeNET^50^ revealed that the 48 consistently down-regulated genes in the individuals with endometriosis were significantly associated with fertility complications such as early pregnancy loss, miscarriage, and spontaneous abortion (Supplementary Figure 2C, Supplementary Table 4).

We also observed that genes related to retinoic acid synthesis and metabolism were persistently decreased in the endometriosis group compared to the normal group during decidualization (Supplementary Table 4). For example, the retinoic acid receptor responder 1 (*RARRES1*) was significantly decreased in endometriosis across all the time points. Aldehyde dehydrogenase 1 family member B1 (*ALDH1B1*) was also significantly decreased in endometriosis relative to normal decreased across all time points. Retinoic acid receptor beta (*RARB*) was decreased on days 2 and 4 of EPC treatment in endometriosis relative to normal. Reprogramming of the endometrium by retinoic acid signaling is critical for endometrial decidualization and early pregnancy^51,52^. Furthermore, altered retinoic acid metabolism also affects endometriotic stromal cell decidualization^53^. Therefore, our results are in line with previous observations, and support the hypothesis that alterations in retinoic acid metabolism drive fertility defects in individuals with endometriosis.

We also observed that the GATA protein binding 6 (*GATA6*) was significantly decreased in the endometriosis group relative to normal cells on days 0, 2, 4, and 6 of EPC treatment. *GATA6* is a *PR* direct target gene^54^ and is upregulated in ectopic endometriotic stromal cells from lesions due to methylation defects^55,56^. Overall, the time course decidualization analysis between normal and endometriosis donors highlighted various pathways that are differentially expressed between the two groups, providing new therapeutic or diagnostic opportunities for endometriosis-associated infertility. Furthermore, our transcriptomic profiling results specifically emphasized the critical roles of the BMP signaling pathways in driving the decidualization processes of the normal endometrium.

### Altered BMP signaling impairs decidualization in the endometrium of individuals with endometriosis

To explore the specific roles of BMP signaling pathways in the decidualization defects observed in individuals with endometriosis, we first assessed the activation of BMP signaling pathways at different time points during time course decidualization. As shown in Figure 3A, in the samples from individuals without endometriosis (n=5), SMAD1/5, the signal transducers of the BMPs, were activated in a time-dependent manner. Upon the EPC stimulation, phosphorylated SMAD1/5 (pSMAD1/5) gradually increased along with the EPC treatment length. However, in the samples derived from individuals with endometriosis (n=4), activation of the BMP signaling pathway was impaired, as manifested by the decreased levels of pSMAD1/5 during the time course EPC stimulation (Figure 3B-C). Gene expression analysis using RT-qPCR demonstrated decidual markers such as *BMP2* and *IGFBP1* showed an increasing trend throughout Day 2 to Day 8 EPC treatment in the stromal cells with blunted induction in the endometriosis cells (Figure 3D-E). These results indicate that while normal decidualizing stromal cells successfully engage BMP/SMAD1/5 signaling, stromal cells from endometriosis failed to show BMP/SMAD1/5 signaling and failed to decidualize.

**Figure 3.**
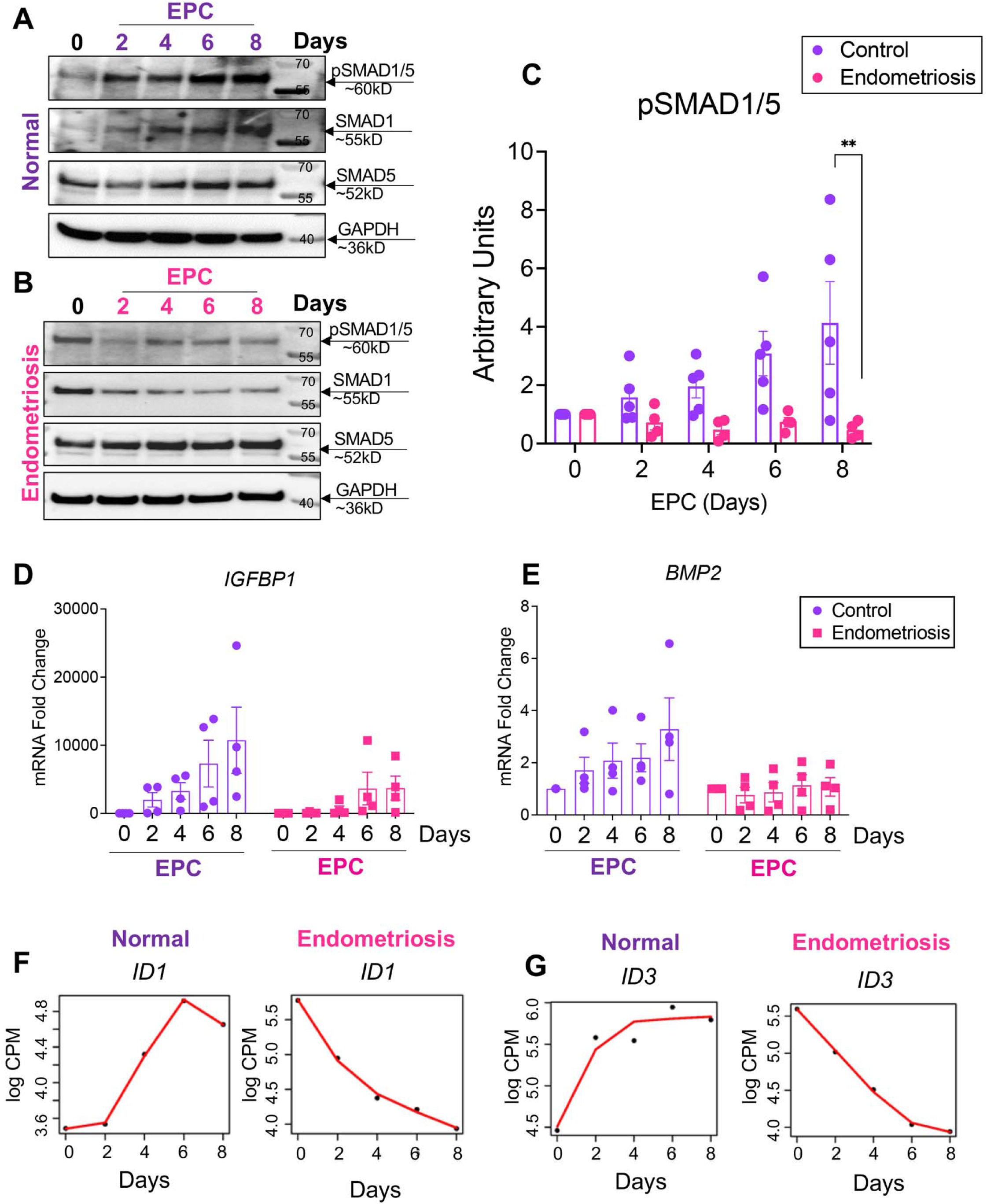
Impaired BMP signaling perturbs decidualization in the endometrium of individuals with endometriosis. A, B) Lysates from endometrial stromal cells of individuals without (A, “normal”) with endometriosis (B) after 2, 4, 6 or 8 days of EPC treatment were probed with antibodies to detect phosphorylated SMAD1/5 (pSMAD1/5), total SMAD1, total SMAD5, or GAPDH expression. C) Densitometric analysis of pSMAD1/5 in the EPC-treated stromal cells from individuals without (n=5) or with endometriosis (n=4). D,E) Quantitative reverse transcriptase PCR (qRT-PCR) was used to determine the expression of *BMP2* and *IGFBP1* in the endometrial stromal cells from individuals without (C, n=4) or with endometriosis (D, n=4). F,G) Time course analysis from the RNAseq analysis comparing the increasing gene expression patterns of *ID1* and *ID3* in normal and decreasing gene expression pattern in endometriosis stromal cells. Histograms (C,D,E) represent mean +/- standard error of the mean analyzed using a One Way ANOVA with Tukey’s posttest for multiple comparisons (C). *EPC,* 35nM estradiol, 1µM medroxyprogesterone acetate, 50µM cAMP.

The discrepancy in response to the EPC treatment observed between individuals with and without endometriosis was also identified from the RNA-seq data. Figure 3F-G shows exemplary genes that present contrasting trends during our time course decidualization analysis. Similar to *BMP2,* the *ID1* and *ID3* genes were progressively increased over the time course treatment in the samples derived from donors without endometriosis while they were conversely down regulated in the endometriosis cohort. Inhibitor of DNA-binding (ID) genes are not only known downstream BMP responsive genes^57,58^ but also are important for endometrial remodeling and decidua formation^59–61^. Such inverted trends substantiated the dysfunctional BMP signaling pathways in the endometriosis groups. Our data indicated that in individuals with endometriosis, an impaired BMP signaling pathway is accompanied by dysfunctional pathways in controlling endometrial transformation.

### Genome-wide binding of SMAD4 reveals differential binding patterns in the endometrium of individuals with and without endometriosis

Upon ligand binding, canonical BMP signaling pathways use SMAD1/5/4 proteins to initiate nuclear transcriptional control. SMAD1/5 forms heterodimers and translocate into the nucleus together with common SMAD4. Our goal was to investigate the mechanisms that underpin defective BMP signaling in individuals with endometriosis during decidualization at the transcriptional level. To do so, we utilized the Cleavage Under Targets & Release Using Nuclease (CUT&RUN) method to profile the genome-wide SMAD4 binding sites in the EPC-treated endometrial stromal cells derived from both individuals with and without endometriosis. We observed distinct pattering of SMAD4 binding activities between the two groups (Supplementary Figure 3A-B). We exemplified the binding activities by showing the Integrative Genomics Viewer (IGV) track view of the *ID1* and *ID3* loci. SMAD4 binding was diminished in the endometriosis groups (Figure 4A). In total, we identified 2060 peaks showing differences in signal intensity between normal and endometriosis groups. Among these, 1190 peaks were less enriched in the endometriosis group, while 870 peaks are more enriched in the endometriosis group. Peak annotation revealed that the majority of these peaks were located within the ± 3 kb promoter region (72.29%) (Figure 4B, Supplementary Table 5). Additionally, we performed the Reactome pathway enrichment^62^ for the genes that were differentially bound by SMAD4 in the endometriosis group (Figure 4C). We found that categories related to ‘signaling by TGFβ family members’, ‘signaling by TGFβ receptor complexes’, and ‘TGFβ activated SMADs’ were enriched, in agreement with our previous transcriptomic results.

**Figure 4.**
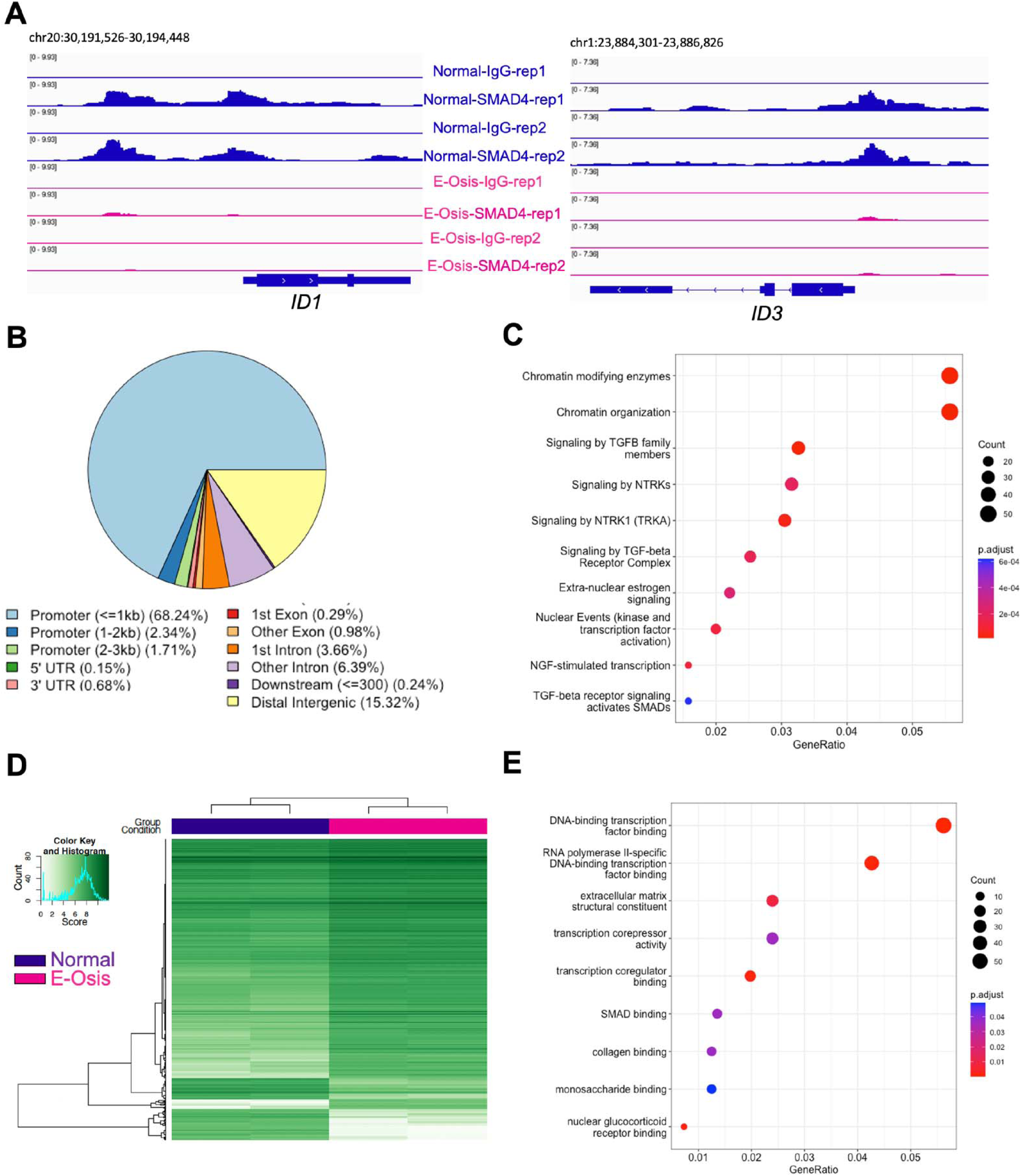
SMAD4 and H3K27ac CUT&RUN reveals differential binding events in the endometrial stromal cells of individuals with endometriosis. CUT&RUN was performed for SMAD4 and H3K27ac in endometrial stromal cells from individuals with or without endometriosis to identify differences or similarities in their genome-wide distribution. A) Genome track views for the *ID1* and *ID3* genes displaying the enriched SMAD4 peaks obtained from the normal cells (blue) that are decreased in the endometriosis cells (pink). B) Peak annotation of the SMAD4 peaks in 2060 peaks showing differences in signal intensity between normal and endometriosis samples, showing many of the differential peaks (72.29%) were located proximal to the promoter region (within ± 3 kb promoter region). C) Reactome analysis showing classification of genes that were differentially bound by SMAD4 in the endometriosis samples. Chromatin modifications and signaling by TGFβ family members were in the top three categories. D) H3K27ac CUT&RUN was performed in endometrial stromal cells from individuals without endometriosis (“Normal”) or with endometriosis (“E-Osis”) after 4 days of EPC treatment. The heatmap shows the peak signal obtained for H3K27Ac in normal versus endometriosis stromal cells. E) Gene ontology classification of the 1122 peaks that were more enriched in the endometriosis samples.

Interestingly, apart from TGFβ related categories, pathways involved in ‘chromatin modifying enzymes’, ‘signaling by NTRKs,’ and ‘NGF-stimulated transcription,’ were also among the top enriched categories (Figure 4C). Neurotrophic tyrosine receptor kinases (NTRKs) are well-documented for their roles in pain and inflammation in endometriosis and are elevated in the endometriotic lesions of affected patients^63–65^. Nerve growth factors^66^ signal through the NTRKs and were recently shown to be associated with endometriosis through genome wide association studies^67^. Hence, our studies suggest that abnormal NTRK signaling may also impact the eutopic endometrium and affect receptivity in patients with endometriosis.

To further delineate the chromatin level differences between the normal and endometriosis groups, we profiled the depositions of histone mark H3K27 acetylation (H3K27ac) in the EPC-treated stromal cells (Supplementary Table 6). H3K27ac modification on the chromatin has been well-defined in the enhancer and promoter regions and is usually accompanied by active transcription activities^68–70^. Similar to SMAD4 binding patterns, H3K27ac marks also showed distinct patterns between normal and endometriosis groups (Figure 4D, Supplementary Figure 4A). We identified 1439 peaks that were more enriched in the normal group and 1122 peaks that were more enriched in the endometriosis group.

For the genes that preferentially have more H3K27ac peaks in the endometriosis group, Gene Ontology (GO) enrichment analysis indicated positive regulation of cell adhesion being the most enriched category, which has been suggested to facilitate lesion establishment at ectopic sites^71^ (Supplementary Figure 4B). GO enrichment on the genes that have less H3K27ac peaks in the endometriosis group indicated that categories involving transcription factor binding, extracellular matrix structural constituent and transcription corepressor activity were deficient during the decidualization process in the endometrium of individuals with endometriosis (Figure 4E). Additionally, genes in the SMAD binding category also have fewer H3K27ac peaks in the endometrial stromal cells from individuals with endometriosis, in agreement with previous transcriptomic results, signifying the dysfunctional BMP signaling pathways in the endometriosis group during decidualization (Figure 4E). Specifically, we showcased the difference in H3K27ac deposition in known progesterone-responsive genes *RARB* and *CEBPA* loci^72,73^ (Supplementary Figure 4C). These results show that defective endometrial transcriptional responses driven by BMP signaling in individuals with endometriosis are detected at the chromatin level, as evidenced by different genome wide SMAD4 and H3K27ac binding patterns in normal versus endometriosis groups.

### Silencing of SMAD1 and SMAD5 perturbs endometrial stromal cell decidualization

To functionally examine the role of BMP signaling pathway in mediating decidualization, we perturbed the SMAD1/5 complex using small interfering RNA (siRNA) in endometrial stromal cells from individuals without endometriosis and treated them with EPC to induce *in vitro* decidualization (Supplementary Figure 5A). The knockdown effect was validated at the transcript and protein level (Figure 5A, Supplementary Figure 5B, Supplementary Table 7). Upon the knockdown of SMAD1/5, we observed that canonical decidualization markers such as *IGFBP1* and *WNT4* were significantly down regulated (Figure 5A). KEGG pathway enrichment on the differentially expressed genes revealed that the TGFβ and FOXO signaling pathways were also enriched within the downregulated group of genes (Figure 5B). FOXO family plays a critical role in regulating progesterone-dependent differentiation and decidualization^74^ and is indispensable for implantation and decidua formation^75^. We highlighted several key genes changes in the heatmap format to visualize the effect of SMAD1/5 perturbation (Figure 5C).

**Figure 5.**
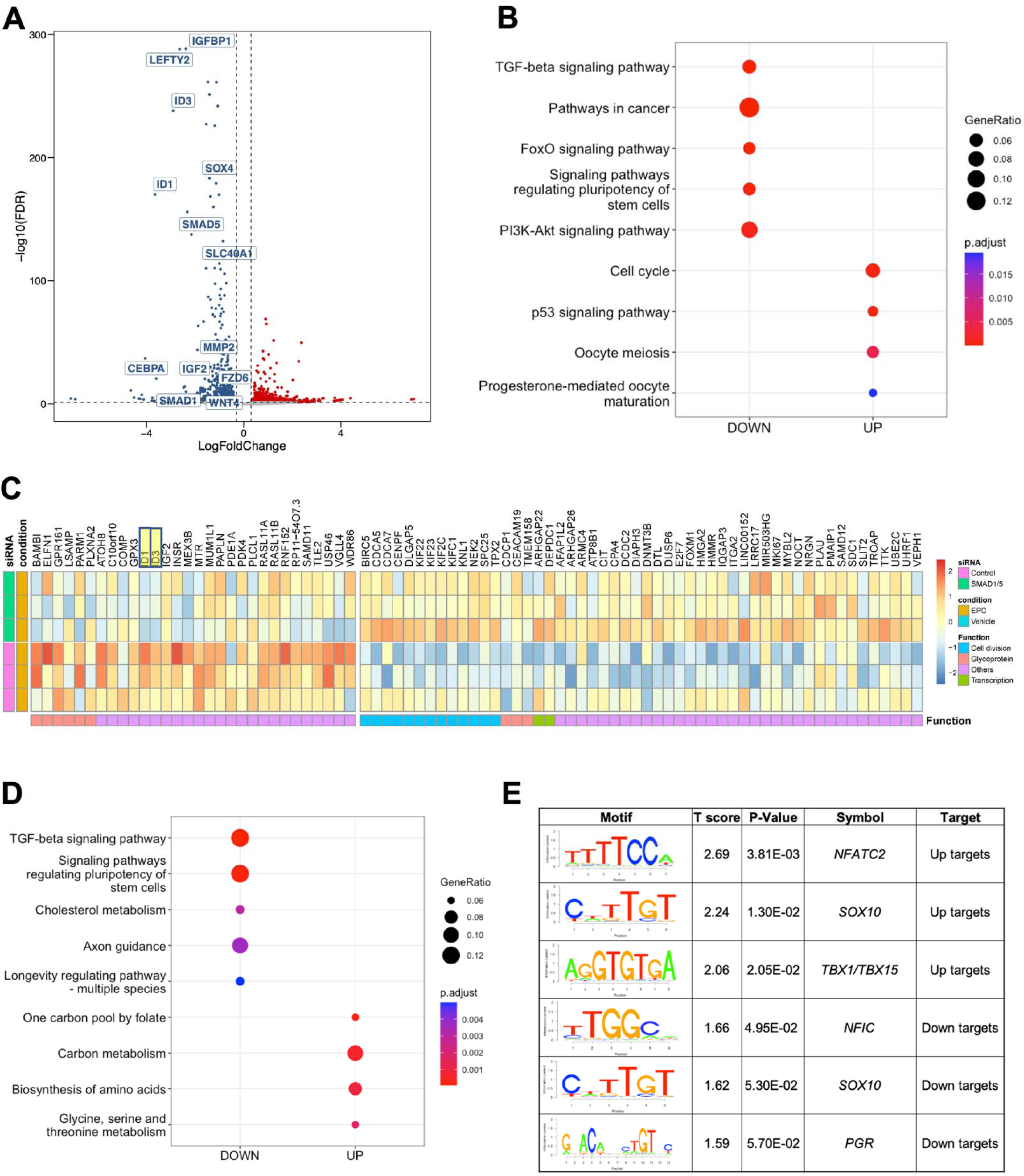
Knockdown of SMAD1 and SMAD5 perturbs decidualization in endometrial stromal cells. A) Volcano plot showing the expression of differentially expressed genes in siCTL + EPC vs. siSMAD1/5 + EPC treated endometrial stromal cells (using a cutoff of Log 2 FC >0.30, < -0.30, FDR < 0.05). Blue indicates genes that are down-regulated in siSMAD1/5 + EPC vs. siCTL + EPC, red indicates genes that are increased. (n=1 individual without endometriosis). B) Dot plot showing the enrichment of genes in key signaling pathways after SMAD1/5 knockdown. C) Heatmap showing the expression and functional classification of key genes following SMAD1/5 knockdown + EPC versus siCTL + EPC treatment (n=3 individuals without endometriosis). D) Binding and expression target analysis (BETA) was used to integrate SMAD4 binding peaks with the transcriptional changes after SMAD1/5 knockdown in EPC-treated endometrial stromal cells. Dotplot displays the gene ontology classification of genes that were activated by SMAD1/5 (i.e., were downregulated by SMAD1/5 and have a SMAD4 binding site). E) Motif analysis was performed on the group genes identified to be SMAD1/5/4 direct targets and displayed as “uptargets” (genes that were increased after SMAD1/5 knockdown and had a SMAD4 peak) or as “downtargets” (genes that were downregulated after SMAD1/5 knockdown and had a SMAD4 peak).

To further map the direct target genes and potential co-factors of SMAD1/5 during decidualization, we used Binding and expression target analysis (BETA)^76^ to consolidate our genomic profiling of SMAD4 and the transcriptomic profiling of SMAD1/5 perturbation. Among the direct targets that were activated by SMAD1/5 (which were down regulated upon SMAD1/5 perturbation and were bound by SMAD4, labeled as Down-targets), were the TGFβ signaling pathway and pathways regulating the pluripotency of stem cells (Figure 5D). We also performed motif analysis on the direct target genes to provide mechanistic insight to the SMAD1/5 mediated gene expression during decidualization. We uncovered potential SMAD1/5 co-repressors such as *NFATC2* and T-box family (*TBX1/TBX15*). *NFATC2* is involved in cGMP-PKG signaling pathways and has a role in regulating immune, inflammatory responses^77^, it is also reported to be elevated in the thin-endometrium patients who usually have deficient implantation and lower pregnancy rate^78^. Acting mainly as repressors, TBX family genes are crucial for embryonic development and tissue differentiation and formation^79^. A recent study has shown that TBX15 was elevated in patients with adenomyosis^80^. As for transcriptional co-activators, apart from canonical pan-tissue co-activator *NFIC*^81^, the motif for the *PR* was enriched in the SMAD1/5 direct target genes, confirming our previous finding that SMAD1/5 may regulate progesterone-responsive genes at the transcriptional level (Figure 5E). We also identified two novel SMAD1/5-activating targets in our analysis, *MALAT1* and *HDAC4.* The direct SMAD4 binding activities in *MALAT1* and *HDAC4* loci were visualized as genome track views in Supplementary Figure 5C-D. *MALAT1* and *HDAC4* are both deeply involved in facilitating decidualization and pathogenesis of endometriosis^82–87^. Our results not only validated the indispensable roles of SMAD1/5 during human decidualization, but also provided additional layers of regulation in the downstream networks of SMAD1/5 mediated BMP signaling pathways.

### BMP2 supplementation enhances the decidualization potential in stromal cells and endometrial assembloids of individuals with endometriosis

To test whether the addition of recombinant BMP2 supplementation could restore the decidual response in the endometrium from individuals with endometriosis, we added BMP2 to endometrial stromal cell cultures and to 3-dimensional endometrial stromal/epithelial co-cultures, or “assembloids” (Figure 6). For the stromal cell experiments, cells derived from the eutopic endometrium of patients with endometriosis were treated as shown in Figure 6A with the EPC cocktail or EPC + BMP2 for a total of 4 days. The extent of decidualization was examined using qPCR analysis of the decidual markers, *FOXO1, PRL, SPP1* and *IGFBP1* (Figure 6B-E)*. FOXO1, PRL* and IGFBP1 all showed an increased trend in expression following EPC+BMP2 treatment versus EPC treatment alone. However, only the expression of *SPP1* changed significantly after BMP2 supplementation (Figure 6B-E). Correspondingly, we observed that the combined addition of EPC + BMP2 synergized the expression of pSMAD1/5 relative to BMP2 or EPC treatment alone (Figure 6F-G).

**Figure 6.**
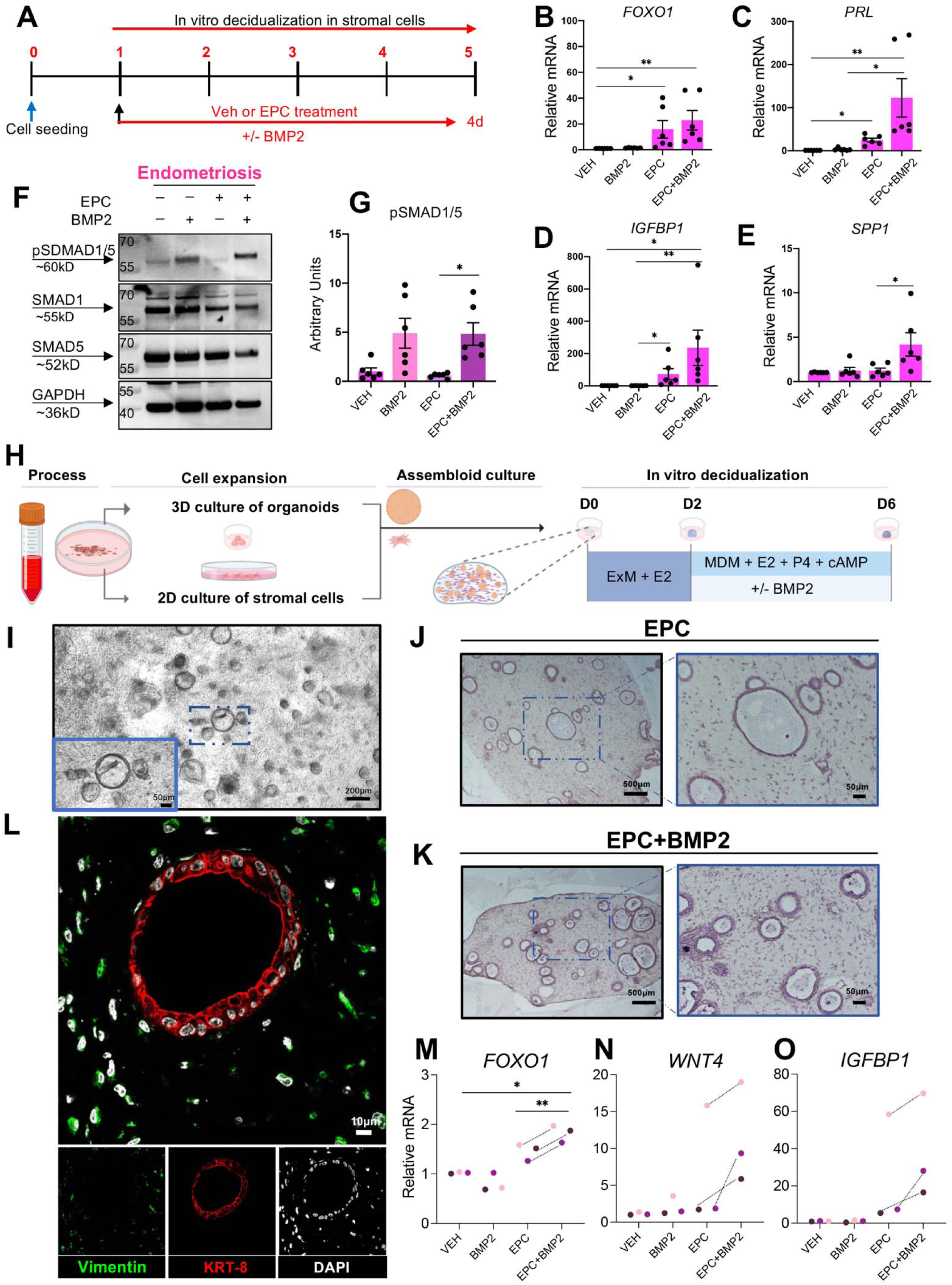
BMP2 supplementation improves the decidualization potential of 2D and 3D endometriosis patient-derived endometrial cultures. A) Experimental outline showing the treatment groups used to test how the addition of recombinant BMP2 affects decidualization in EPC-treated stromal cells from individuals with endometriosis. B-E) qRT-PCR quantification of decidualization markers *FOXO1* (B)*, PRL* (C)*, IGFBP1* (D), and *SPP1* (E) following Vehicle, BMP2, EPC, or EPC + BMP2 treatment in stromal cells from individuals with endometriosis (n=6). F-G) Western blot analysis (F) and quantification (G) of endometrial stromal cells from individuals with endometriosis following 4 days of treatment with Vehicle, BMP2, EPC, or EPC + BMP2. H) Diagram showing the experimental procedure for establishing endometrial epithelial and stromal co-cultures or “assembloids” from endometrial tissues of individuals with endometriosis. After the assembloids were established, they were pre-treated with 10nM estradiol (E2) followed by decidualization with the EPC decidualization cocktail (1 µM MPA, 0.5 mM cAMP and 1 µM E2) +/- 25 ng/ml BMP2 for an additional 4 days. I) Phase contrast micrograph of the endometrial epithelial and stromal assembloids showing the endometrial epithelial organoids and the distribution of stromal cells in the collagen matrix. J-L) Histological analysis of cross sections obtained from the endometrial assembloids stained with hematoxylin and eosin (J, K) or using immunofluorescence using vimentin (green), cytokeratin 8 (KRT-8, red) or DAPI (white) (L). M-O) qRT-PCR analysis of decidualization markers, *FOXO1* (M), *WNT4* (N), or *IGFBP1* (O) in the endometrial assembloids treated with Vehicle, BMP2, EPC, or EPC + BMP2. Histograms represent mean +/- standard error of the mean. Histograms were analyzed using a one-way ANOVA with a posthoc test, *<0.05, **<0.001.

To test the impact of BMP2 supplementation on the decidualization potential of endometrial stromal and epithelial assembloids we generated co-cultures as previously described using the strategy outlined in Figure 6H^88^. Individual cultures of endometrial stromal cells and epithelial organoids were established. Four days after initial establishment, the co-cultures were created by encapsulating stromal cells and epithelial organoids in the collagen matrix cultured in expansion medium (ExM) supplement with E2 for 2 days in a shaking system. The culturing medium was then switched to a minimal decidualization media (MDM) containing the decidualization cocktail (EPC) +/- BMP2 for an additional 4 days. Live assembloid cultures were visualized using phase microscopy (Figure 6I) and using histology or fluorescence microscopy after fixation and staining (Figure 6J-L). qPCR analysis of the treated assembloids showed that the BMP2 + EPC supplementation significantly increased *FOXO1* expression and caused an increased trend in the expression of decidual markers *WNT4* and *IGFBP1* relative to EPC treatment alone (Figure 6M-O). We also observed fewer FOXJ1-positive ciliated cells in the endometrial assembloids following EPC + BMP2 treatment, compared with EPC treatment alone (Supplementary Figure 6). These results confirm that BMP2 supplementation increases decidual gene expression in endometrial stromal or in 3D assembloid cultures from individuals with endometriosis.

## METHODS

### Ethics statement and endometrial sample collection

All patient specimens were collected following informed patient consent approved under protocol H-21138 and through the Human Tissue and Pathology Core at Baylor College of Medicine (BCM), following guidelines approved by the Institutional Review Board at BCM. Samples are maintained using de-identified codes to preserve confidentiality. Endometrial samples were obtained from women with confirmed endometriosis (n= 7, mean age, 36.7 +/- 6.9) or from women without endometriosis (n= 7, mean age, 38.4 +/- 5.3) undergoing endometrial biopsies or hysterectomies. Samples categorized in the normal group were free of endometriosis, according to pathology examination reports.

### Establishment and decidualization of primary endometrial stromal cells

Primary endometrial stromal cells were isolated from surgically resected endometrial biopsies, which were immediately placed in stromal culturing media, DMEM/F12 (Gibco #11330032) supplemented with 10% FBS, 1% Antibiotic-Antimycotic (Gibco #15240062), and 100 µg/mL of Primocin (InvivoGen, Cat # MSPP-ANTPM2). Endometrial biopsies were cut into small pieces, digested in Hanks’ Balanced Salt Solution (HBSS) containing 5mg/mL of collagenase (Sigma, #C0130-1G) and 0.2mg/mL of DNase I (Sigma, Cat #DN25-100MG), and then incubated at 37°C for 20 minutes on an orbital shaker at 120 rpm. After incubation, the digested tissues were spun down, and the pellets were resuspended in stromal culturing media. The cell suspension was passed through 100 µm cell strainers and then 20 µm cell strainers. The cell fraction from the flowthrough after the 20 µm cell strainers contains the stromal cells and was cultured in stromal culturing media. Stromal cells were passaged once they reached 90% confluency. For decidualization, stromal cells were seeded on 12-well plates at 2×10^5^ cells/well and 10-cm dishes at 1×10^6^ cells/dish and treated with phenol red-free DMEM/F12 (Gibco #11039021) supplemented with 2% charcoal-stripped FBS and EPC cocktail (1 µM MPA, Sigma Cat#1378001- 200MG, 0.05 mM cAMP, Axxora Cat #JBS-NU-1502-50, and 35 nM E2, Sigma Cat #E1024-1G).

### Establishment and decidualization of endometrial assembloids

Establishment of endometrial assembloids was performed following a previously published protocol^88^ with minor modifications. In brief, primary endometrial stromal cells and glandular epithelial organoids were established from human endometrial samples. After culturing epithelial organoids and stromal cells separately for two passages, the two were mixed gently at a ratio of 1:2 (v/v) and resuspended in 20 times ice-cold Collagen (Sigma, Cat # C0130-1G). Cells were then aliquoted in 20 µl volumes into a 48-well plate and allowed to polymerize at 37□°C for 45 min, after which the collagen assembloid droplets were overlayed with 500µl of Expansion Medium and maintained in a 37 °C cell culture incubator. The collagen droplets were maintained under constant shaking at 90 rpm for 48 hours. (Expansion Medium consists of: Advanced DMEM/F12 (Invitrogen, Cat #12634010), supplemented with 1X N2 supplement (Invitrogen, Cat # 17502048), 1X B27 supplement (Invitrogen, Cat # 12587010), 100 μg/ml Primocin (Invivogen, Cat # MSPP-ANTPM2), 2 mM L-glutamine (Invitrogen, Cat # 25030-024), 500 nM A83-01 (Sigma, Cat #2939), 10% R-Spondin conditioned media, 10 mM Nicotinamide (Sigma, Cat # N0636-100G), 1.25 mM N-acetyl-L-cysteine (Sigma, Cat # A9165-5G), 10% Noggin conditioned media, 10% WNT3a conditioned media, 100 ng/ml FGF10 (Peprotech, Cat # 100-26), 50 ng/ml HGF (Peprotech, Cat #100-39), 50 ng/ml EGF (Peprotech, Cat # AF-100- 15) and 10nM E2 (Sigma, Cat. #E1024-1G). Conditioned media was produced in HEK293 cells and obtained from the Center for Digestive Diseases and Organoid Core Facility at Baylor College of Medicine. To induce decidualization of the assembloids, the culturing media was changed to decidualization media in minimal differentiation media (MDM) (Advanced DMEM/F12 supplement with 1X N2 supplement, 1X B27 supplement, 100 μg/ml Primocin, 2 mM L-glutamine, 1 µM MPA, 0.5 mM cAMP and 1 µM E2) supplemented with or without rhBMP2 (R&D, Cat #355-BM-010/CF) at 25ng/ml. Media was refreshed every 48 hours.

### Histological assessment of endometrial assembloids

Assembloids were fixed in 4% PFA at room temperature for 15 mins and then immobilized in the Histogel (Thermo Fisher, Cat #22-110-678). After processing the assembloids in Histogel, assembloid blocks were dehydrated through a series of ethanol washes and processed for paraffin embedding at the Human Tissue Acquisition and Pathology Core at Baylor College of Medicine. Paraffin blocks were sectioned using 5 µm thick sections. Assembloid sections were deparaffinized in Histoclear and rehydrated in a series of 100%, 95%, 80%, and 70% ethanol washes, followed by washing in dH20. For identifying the morphological structures, assembloid sections were sectioned and stained with hematoxylin and eosin. For immunofluorescence staining, assembloid sections were heated in boiling 10mM sodium citrate, pH 6.0 for 20 minutes for antigen retrieval and quenched in 3% hydrogen peroxide for 10 minutes. After blocking with 3% BSA for 1 hour, assembloid sections were incubated with primary and fluorescent secondary antibodies according to the manufacturer’s instructions and nuclei were stained with 1mg/ml DAPI (1:100- dilution, ThermoFisher, Cat # D1306). The stained slides were mounted in VECTASHIELD antifade mounting medium (Vector Laboratories #H-1000-10). Fluorescence images were taken on a Zeiss LSM780 confocal microscope at the Optical and Vital Microscopy Core at Baylor College of Medicine.

### RNA extraction and quantitative PCR from endometrial stromal cells and endometrial assembloids

The RNAs of endometrial stromal cells were extracted by using QIAGEN RNeasy micro kit (QIAGEN, Cat #74004) according to the manufacturer’s instruction. The assembloids were lysed in Trizol reagent (Life Tech, Cat #10296010) and the RNAs were extracted by using Direct-zol RNA microprep kit (Zymo Research, Cat #R2062). A total of 50-200ng of RNA from each sample was transcribed into cDNA by using qScript cDNA supermix (Quantabio, Cat #95048-100). Real time quantitative PCR was performed on Bio-Rad CFX384 Touch Real-Time PCR Detection System. Fold changes of target genes were calculated using delta delta Ct method and normalized *GAPDH*^89^. Primer sequences are listed in Supplementary Table 8.

### Protein extraction and western blotting

Cells were washed with ice-cold 1×DPBS and lysed in M-PER mammalian protein extraction reagent (ThermoFisher, Prod#78505) supplement with protease inhibitor cocktail (ThermoFisher, Cat #78437) and phosphatase inhibitor cocktail (ThermoFisher, Cat #78426). Protein concentrations were quantified by using Pierce BCA protein assay kit (ThermoFisher, Cat #23225). A total of 20 µg of protein lysate was loaded onto 4- 12% Bis-Tris Plus Mini protein gels (ThermoFisher, Cat # NW04122BOX) and transferred onto nitrocellulose membranes (Bio-Rad, Cat #1704270). The membranes were blocked with 5% non-fat milk in TBST buffer for an hour at room temperature and then incubated with primary antibodies at 4 °C overnight. Antibody information is listed in Supplementary Table 9. The next day, the membranes were probed with HRP-conjugated secondary antibodies (Jackson ImmunoResearch) for two hours at room temperature and protein bands were visualized by using SuperSignal West chemiluminescent substrate (Pierce) on Bio-Rad Chemidoc Touch Imaging system. Protein bands were quantified by using Image Lab software (Rio-Rad).

### Gene expression profiling using RNA sequencing

All sequencing data are available in the NCBI Gene Expression Ominibus under SuperSeries GSE243158. The secure token for reviewer access is wfolcuqovnwxnkd.

#### Time course EPC studies

Endometrial stromal cells from 4 normal and 4 endometriosis samples were treated with 35nM estradiol, 1µM medroxyprogesterone acetate and 1µM 8-Br-cyclic AMP for 0, 2, 4, 6, or 8 days. RNA expression profiles were obtained at each timepoint using RNA sequencing analyses (20-30 million paired-end reads using NovaSeq System from Novogene Corporation Inc. Reads were trimmed with fastp v0.23.2 and aligned using STAR 2.7.10a to human genome assembly GRCh38.p13. Differentially expressed genes between normal patients and endometriosis EPC-treated cells were obtained by comparing to the baseline samples (Day 0). Significantly changed genes during the time course treatment were obtained using an ANOVA F-test using an FDR <0.05 from patients without (n=3) and with endometriosis (n=4). All DEGs are presented in Supplementary Table 1. EnrichR^90–92^ was used to identify the gene ontology classifications (Supplementary Table 2), as well as ENCODE and ChEA consensus transcription factors known to regulate differentially expressed genes in the normal and endometriosis EPC-treated stromal cells (Supplementary Table 3). Differentially expressed genes between normal and endometriosis stromal cells were obtained by comparing transcripts at each time point of treatment. Differentially expressed genes between normal (n=4) and endometriosis (n=3) were identified using a Wald test with a cutoff values of fold-change > 2 or < 1/2 and FDR < 0.05 (Supplementary Table 4).

#### SMAD1/5 siRNA studies

Endometrial stromal cells from 3 individuals without endometriosis were treated with either 80nM negative control (siCTL, Horizon Cat #D-001810-10-20) or 40nM of SMAD1 plus 40 nM of SMAD5 (siSMAD1/5, Horizon Cat # L-012723-00-0005 & L-015791-00-0005) siRNA followed by 4 days’ EPC treatment. Cells were transfected using Lipofectamine RNAiMAX (LifeTechnologies, Cat #13778500). RNAs were isolated by using QIAGEN RNeasy micro kit and subjected to RNA sequencing analysis to identify differentially regulated transcripts. Samples were normalized through effective library sizes and DESeq2 was used identify differentially expressed genes between the siCTL and siSMAD1/5 EPC-treated cells using an FDR < 0.05 (Supplementary Table 7).

### SMAD4 and H3K27 genome-wide binding studies using CUT & RUN

CUT&RUN experiments were performed following a previously published protocol^93^. After endometrial stromal cells were treated with EPC for 4 days, they were collected by digesting with 0.25% Trypsin (ThermoFisher, Cat #25200056) for 3 min. After the digestion, cells were pelleted down at 300 x g for 3 min and viably frozen down in the freezing medium (90% FBS with 10% DMSO) until experiment day. On the day of the experiment, cell vials were quickly thawed and washed 3 times with washing buffer (20 mM HEPES pH 7.5, 150 mM NaCl, 0.5 mM Spermidine, 1 X Protease Inhibitor). For each reaction, 1.3 x 10^6^ cells were used for the subsequent Concanavalin A bead binding step. After 10 min incubation with Concanavalin A beads, bead-cell complexes were resuspended in 100 μl antibody buffer (washing buffer supplemented with 0.01% digitonin, and 2mM EDTA) per reaction. 1 μl of IgG antibody (Sigma, Cat #I5006), H3K27ac (Cell Signaling, Cat #8173) and SMAD4 antibody (Abcam, Cat #ab40759) were added to each reaction respectively. After overnight incubation at 4 °C, bead-cell complexes were washed twice with 200 μl cold dig-washing buffer (washing buffer supplemented with 0.01% digitonin) and resuspended in 50 μl cold dig-washing buffer with 1 μl pAG-MNase (EpiCypher, Cat #15-1016). After incubation at room temperature for 10 min, bead-cell complexes were washed twice with 200 μl cold dig-washing buffer and resuspended in 50 μl cold dig-washing buffer, then 1 μl 100 mM CaCl_2_ was added to each reaction. The mixture was incubated at 4 °C for 2 hours and the reaction was stopped by adding 50 μl stop buffer (340mM NaCl, 20 mM EDTA, 4 mM EGTA, 0.05% Digitonin, 100 ug/mL RNase A, 50 mg/mL glycogen, 0.5 ng E. coli DNA Spike-in (EpiCypher, Cat #18-1401) and incubated at 37 °C for 10 min. The supernatant was collected and subjected to DNA purification with phenol-chloroform and ethanol precipitation. Sequencing libraries were prepared using NEBNext Ultra II DNA Library Prep Kit (New England BioLabs, Cat #E7645) following manufacture’s protocol. Paired-end 150 bp sequencing was performed on a NEXTSeq550 (Illumina) platform and each sample was targeted for 10 million reads. Sequencing raw data were de-multiplexed by bcl2fastq v2.20 with fastqc for quality control and then mapped to reference genome hg19 by Bowtie2, with parameters of --end-to-end --very-sensitive --no-mixed --no-discordant --phred33 -I 10 -X 700. For Spike-in mapping, reads were mapped to E. coli genome U00096.3. Spike-in normalization was achieved through multiply primary genome coverage by scale factor (100000 / fragments mapped to E. coli genome). CUT&RUN peaks were called by Model-based Analysis of ChIP-Seq (MACS/2.0.10)^94^ with the parameters of -f BAMPE -g 1.87e9 -q 0.05 (H3K27ac) or -q 0.1 (SMAD4). Track visualization was done by bedGraphToBigWig20, bigwig files were imported to Integrative Genomics Viewer for visualization. For peak annotation, genomic coordinates were annotated by ChIPseeker^95^. Differential binding analysis and clustering were conducted using DiffBind^96^. Direct targets motif analysis was conducted through Binding and Expression Target Analysis (BETA)^76^ with parameter BETA plus –p –e –k LIM –g hg19 --gs hg19.fa --bl. Annotated peak files were included in Supplemental Table 5 (SMAD4) and Supplemental Table 6 (H3K27ac).

## DISCUSSION

Our study presents transcriptomic evidence supporting the hypothesis that patients with endometriosis display abnormal decidualization programs that can partially explain the elevated infertility rates within that population. By examining the time-dependent endometrial response to hormones in individuals with and without endometriosis, we showed that some key pathways were conserved between the two groups, such as interleukin-2 signaling and TGFβ regulation of extracellular matrix, while others were unique to each group, such BDNF signaling in normal endometrium and cardiomyocyte hypertrophy in endometriosis. Accordingly, inflammatory-related genes that are controlled by IL2 signaling, such as *IL6, IL24,* and *IL1R1*, showed similar trends in the normal and endometriosis groups during the time-course decidualization analyses. A similar case was found for TGFβ-related genes. However, others also showed different activation patterns when comparing endometrial stromal cells from individuals with and without endometriosis. These included *NFE2L1,* which increased in the endometriosis group but decreased in the normal group, and *RGS5,* which decreased more extensively in the normal group when compared to endometriosis. Decidualization induces extensive genetic and epigenetic remodeling programs in the endometrium that result in morphological and functional specialization of the tissue. Our results suggest that while the endometrial stromal cells from individuals with endometriosis engage similar transcriptional responses to those from individuals without, alterations in other key pathways may compromise their complete decidualization potential.

To identify the regulatory factors that could be driving the different transcriptional responses between the endometrial stromal cells from individuals with or without endometriosis, we explored consensus gene targets in the ENCODE and ChEA Transcription Factor Targets datasets^36,37^. This analysis indicated that CEBP/β is a major transcription factor controlling the transcriptional response to decidualization in the normal endometrium, controlling genes such as *FBXO32, YARS,* and *MMP19*. CEBP/β has also been shown to be a master regulator of human endometrial cell decidualization, which controls the expression of *PGR* by directly binding to its promoter^38,39^. Analysis of endometrial stromal cells from individuals with endometriosis identified NFE2L2 and SMAD4 as the top two transcription factors controlling gene expression during decidualization. NFE2L2 is a central factor controlling the intracellular response to stress and was previously shown to be activated in the endometrial epithelial cells of cows exposed to heat stress^97^. NFE2L2 (also known as NRF2) controls the expression of antioxidant genes in the cell by binding to DNA antioxidant response elements (or “AREs”)^98^. One class of genes controlled by NFE2L2 are the glutathione peroxidase genes (i.e., *GPX3* and *GPX4*), which play key roles in the control of cellular oxidative stress damage^47^. Hence, our data supports theories regarding the altered response to oxidative stress in the endometrium of individuals with endometriosis as a possible leading cause for impaired decidualization^99^. Others have also suggested that impaired response to oxidative stress through defective iron metabolism is an underlying factor in women with recurrent pregnancy loss^100^.

The ENCODE and ChEA Transcription Factor gene target analysis also identified SMAD4 as a major regulatory factor controlling transcription in endometriosis. The TGFβ signaling pathway was also notably altered in the endometrium of individuals with endometriosis when we directly compared the genes that were differentially regulated between normal and endometriosis groups at each time point during decidualization. (Figure 2E) For example, the expression of the canonical SMAD1/SMAD5 targets, *ID2* and *ID3,* were significantly decreased during days 4 and 6 of EPC treatment. *GREMLIN2,* which is a secreted antagonist of the BMPs was increased in the endometrial stromal cells from individuals with endometriosis on Day 0. The expression of *FST* was decreased in the endometriosis group after 2, 4, 6 and 8 days of EPC treatment. Thus, using gene ontology analysis and upstream regulatory factor analyses, we concluded that the transcriptional control by BMP/SMAD signaling was a key pathway controlling decidualization the endometrium of individuals without endometriosis that was perturbed in individuals with endometriosis.

To further characterize the genome-wide distribution of the downstream effectors of the BMPs, we used CUT&RUN to detect SMAD4 binding events in stromal cells from individuals with and without endometriosis after EPC treatment. To detect the chromatin-level changes between the two cohorts, we also mapped H3K27ac marks in the endometrial stromal cells from the normal and endometriosis groups. The binding studies showed that there were notable changes in the distribution of both SMAD4 and H3K27ac between the endometrial stromal cells of individuals without and with endometriosis, suggesting that the gene expression changes were a result of altered transcription factor binding events. This was further confirmed by intersecting the SMAD4 binding events with differentially expressed genes following SMAD1/SMAD5 siRNA-mediated knockdown in endometrial stromal cells treated with EPC to induce *in vitro* decidualization. We first identified that the double knockdown of SMAD1 and SMAD5 blunted the decidualization capacity of endometrial stromal cells, as evidenced by the decrease of the canonical decidualization markers, *IGFBP1* and *WNT4*. Merging of the SMAD4 binding peaks and downregulated genes after SMAD1/5 knockdown showed enrichment of genes with consensus sequences related to transcription by NFIX, SOX10, and PR. Progesterone receptor is the master regulator of decidualization^101,102^, suggesting that impaired BMP/SMAD1/5 signaling perturbs transcriptional activation of PR, blunting endometrial cell reprogramming. Furthermore, we identified direct SMAD4 binding sites on the genes of the *MALAT1* and *HDAC4* genes, which are critical for endometrial stromal cell decidualization^82–87^. Our results show molecular evidence that impaired BMP/SMAD signaling underlies the decidualization defects in the endometrium of individuals with endometriosis.

To verify these findings, we tested whether the addition of recombinant human BMP2 to decidualizing cultures of endometrium could increase endometrial decidualization markers in individuals with endometriosis. Previous studies have shown that ectopic expression of BMP2 in endometrial stromal cells could potentiate decidualization in the normal endometrium^103^. We used both 2D endometrial stromal cells as well as 3D epithelial/stromal cocultures or “assembloids” to recapitulate paracrine signaling events. We found that relative to EPC treatment alone, the addition of BMP2 + EPC increased the expression of canonical decidual genes in both the stromal cell and assembloid cultures of endometrium from individuals with endometriosis. Our findings indicate that BMP2 and the downstream activated signaling pathways are defective in the endometrium of patients with endometriosis and that BMP2 supplementation may correct the defect.

BMPs are subgroups of the TGFβ ligand family and BMP signaling pathways are indispensable in the female reproductive tract, especially during early pregnancy establishment^49,104^. Upon BMP ligand binding, the serine-threonine kinase receptors (ALK3/ALK6/ACVR1/BMPR2/ACVR2A/ACVR2B) will subsequently phosphorylate the signal transducers, SMAD1 and SMAD5 and phosphorylated SMAD1/5 will then form homodimers and translocate into the nucleus together with a common SMAD4 protein to initiate transcriptional programming. BMP signaling pathways are key in transforming the maternal endometrium into a receptive environment for further support embryo implantation. From the uterine-specific knockout mouse models, BMP ligands^105,106^, kinase receptors and SMAD signal transducers^38,107–110^ are essential in decidualization and implantation, which are prerequisites for the establishment of a healthy pregnancy. Apart from regulating the decidualization and implantation process, BMP signaling pathways are also involved in immunomodulation in the endometrium. Conditional deletion of *Bmpr2* in the mouse uterus diminishes the uterine natural killer cell populations, which regulate the immune response in the endometrium, preventing the rejection of the embryo as a foreign entity. Such an immune-privileged microenvironment is crucial in the early stages of pregnancy^108^.

Indeed, the essential roles that BMP signaling pathways play in cell differentiation, proliferation and anti-inflammation potentiates its significance in the context of endometriosis. Interestingly, BMP2 levels were decreased in the peritoneal fluid of women with endometriosis^111^. Recent large-scale genome-wide association studies (GWAS) also identified *BMPR2* as one of the endometriosis risk loci^67^. In women with recurrent implantation loss, *BMP7* was identified to harbor a deleterious mutation that was predicted to be disease-causing^112^. However, the intricate relationship between endometriosis and the BMP signaling pathway is still not clearly understood.

Using transcriptomic and genome-wide binding analyses in patient-derived 2D and 3D-endometrial cultures, we show that abnormal BMP signaling pathways may affect fertility in individuals with endometriosis by directly affecting the decidualization process of the endometrium. Our findings presented here corroborate previous studies that noted the abnormal endometrial response to hormones in individuals with endometriosis^28–30^. However, they also reveal alterations in new pathways, such as the BMP/SMAD signaling pathways, oxidative stress responses, and retinoic acid signaling pathways, opening new potential avenues for the development of biomarkers or therapeutics for endometriosis-associated infertility.

## DATA AVAILABILITY STATEMENT

Sequencing data are available in the NCBI Gene Expression Ominibus under SuperSeries GSE243158. The secure token for reviewer access is wfolcuqovnwxnkd.

## ACKNOWLEDGEMENTS

We are grateful to Dr. Martin M. Matzuk (M.M.M) for his support and guidance on this project. Studies were supported by *Eunice Kennedy Shriver* National Institute of Child Health and Human Development grants R00-HD096057 (D.M.), R01-HD105800 (D.M.), R01-HD032067 (M.M.M.) and R01-HD110038 (M.M.M). Diana Monsivais, Ph.D. holds a Next Gen Pregnancy Award (NGP10125) from the Burroughs Wellcome Fund.

## AUTHOR CONTRIBUTIONS

Study conception and design: Z.L., S.T., P.J., T.G., D.M. Performed experiment or data collection: Z.L., S.T., P.J., T.G., T.N.D, J.G., L.A.R., X.G., D.M.. Computation and statistical analysis: Z.L., S.T., P.J., T.G., D.M. Data interpretation and analysis: Z.L., S.T., P.J., T.G., D.M. Writing, reviewing, and editing: All. Supervision: D.M.

## SUPPLEMENTARY INFORMATION

**Supplemental Figure 1.**
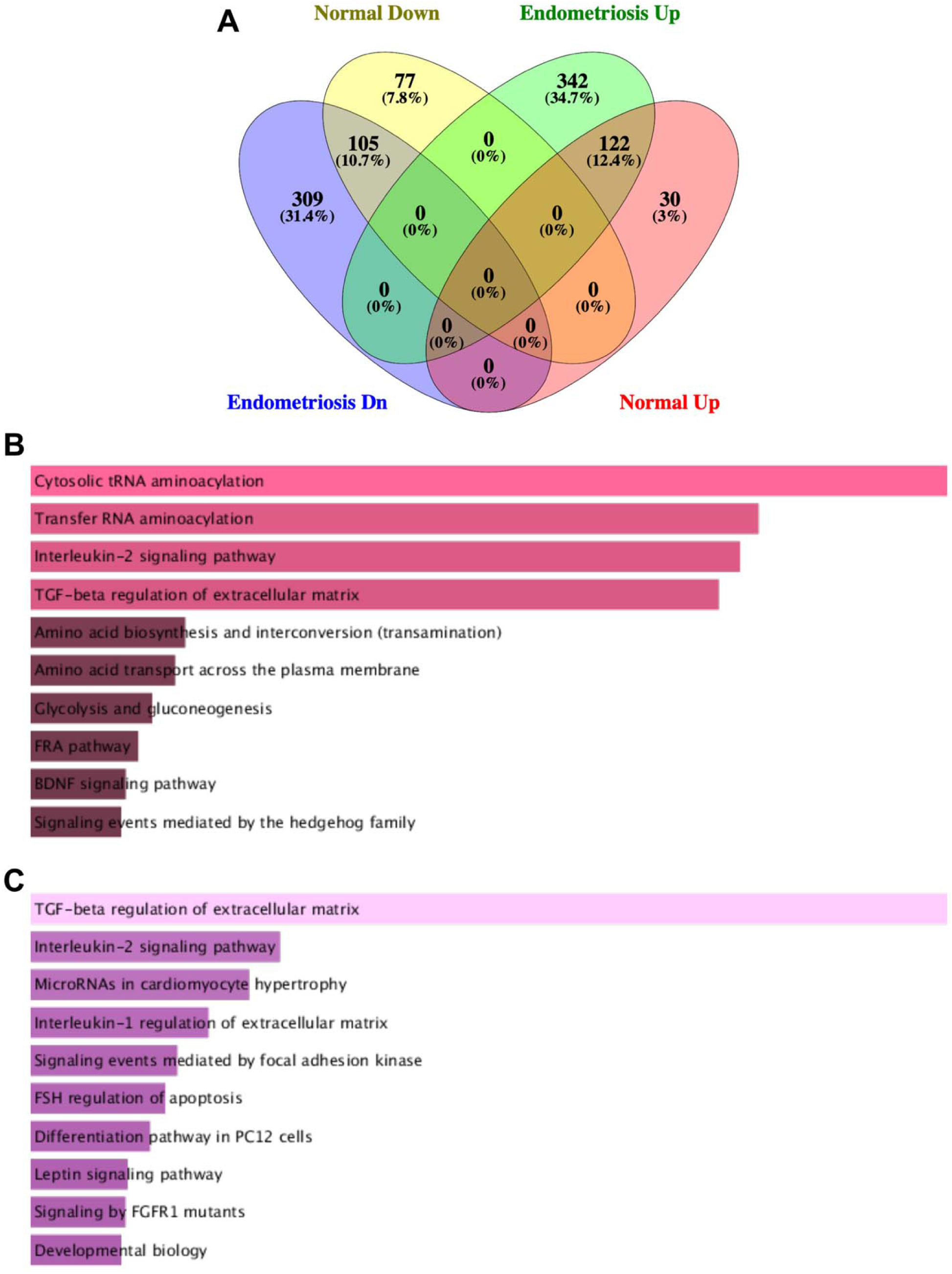
Transcriptomic and gene ontology analysis of the time course decidualization datasets. A) A Venn diagram was used to display the number of conserved genes between the normal and endometriosis groups during the time course decidualization program. Genes that showed a significant change on Day 8 of decidualization were used (>1.4, <0.4-fold change, FDR < 0.05). B-C) Gene ontology analysis of the genes that showed a significant change (>1.4, <0.4-fold change, FDR < 0.05) on Day 8 of decidualization in the normal (B) or endometriosis (C) groups.

**Supplemental Figure 2.**
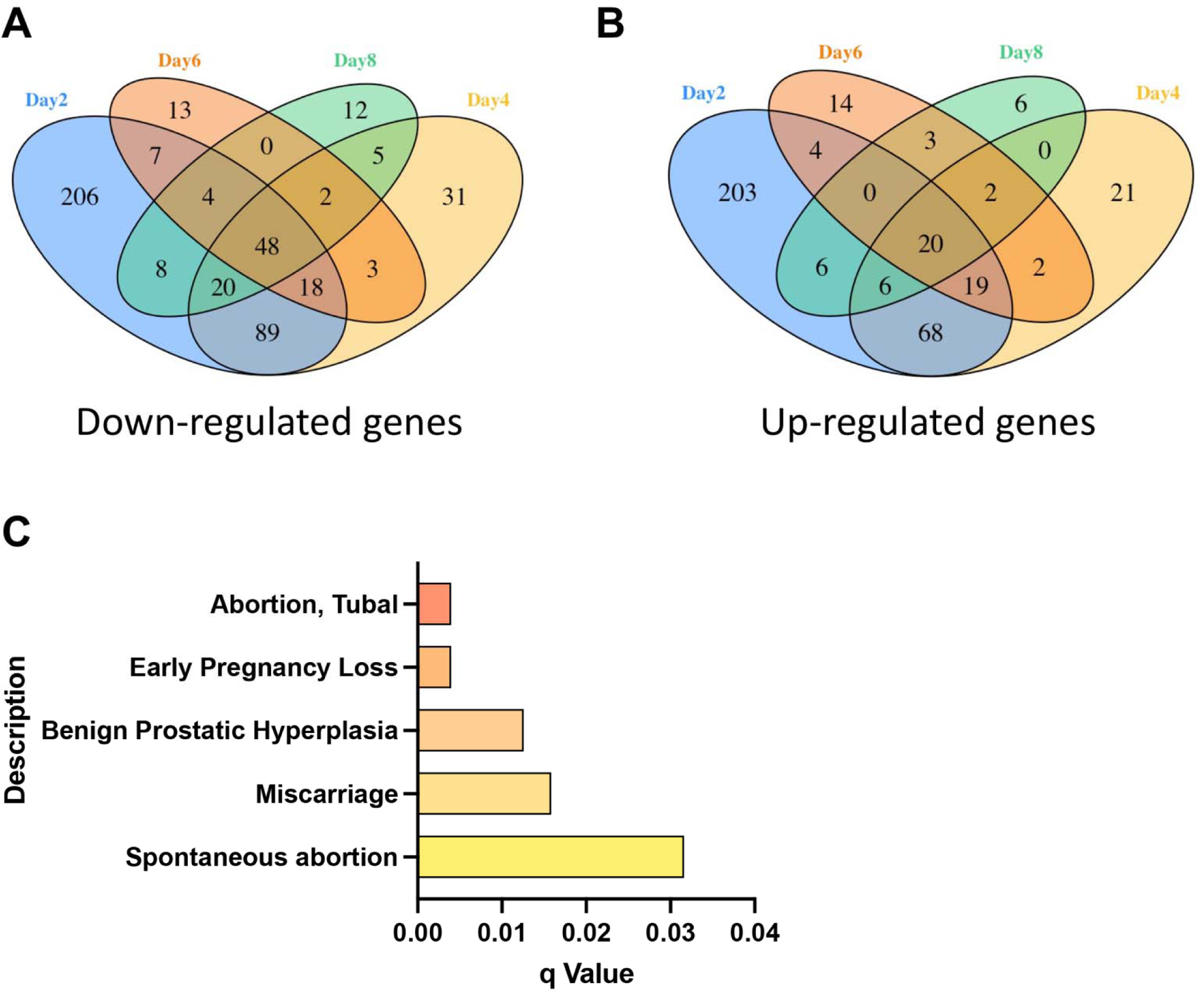
Transcriptomic and classification of genes involved in the decidualization of endometrial stromal cells from individuals with and without endometriosis. A-B) A Venn diagram was used to display the number of conserved downregulated (A) and upregulated (B) genes in the time course decidualization of endometrial stromal cells from individuals with endometriosis. The analysis identified that 48 genes were consistently down-regulated (A) and 20 genes were consistently up-regulated (B) regardless of the EPC treatment length. C) DisGeNET analysis for the 48 genes that were consistently down regulated in the stromal cells from individuals with endometriosis relative to individuals without endometriosis during the time course decidualization treatment.

**Supplemental Figure 3.**
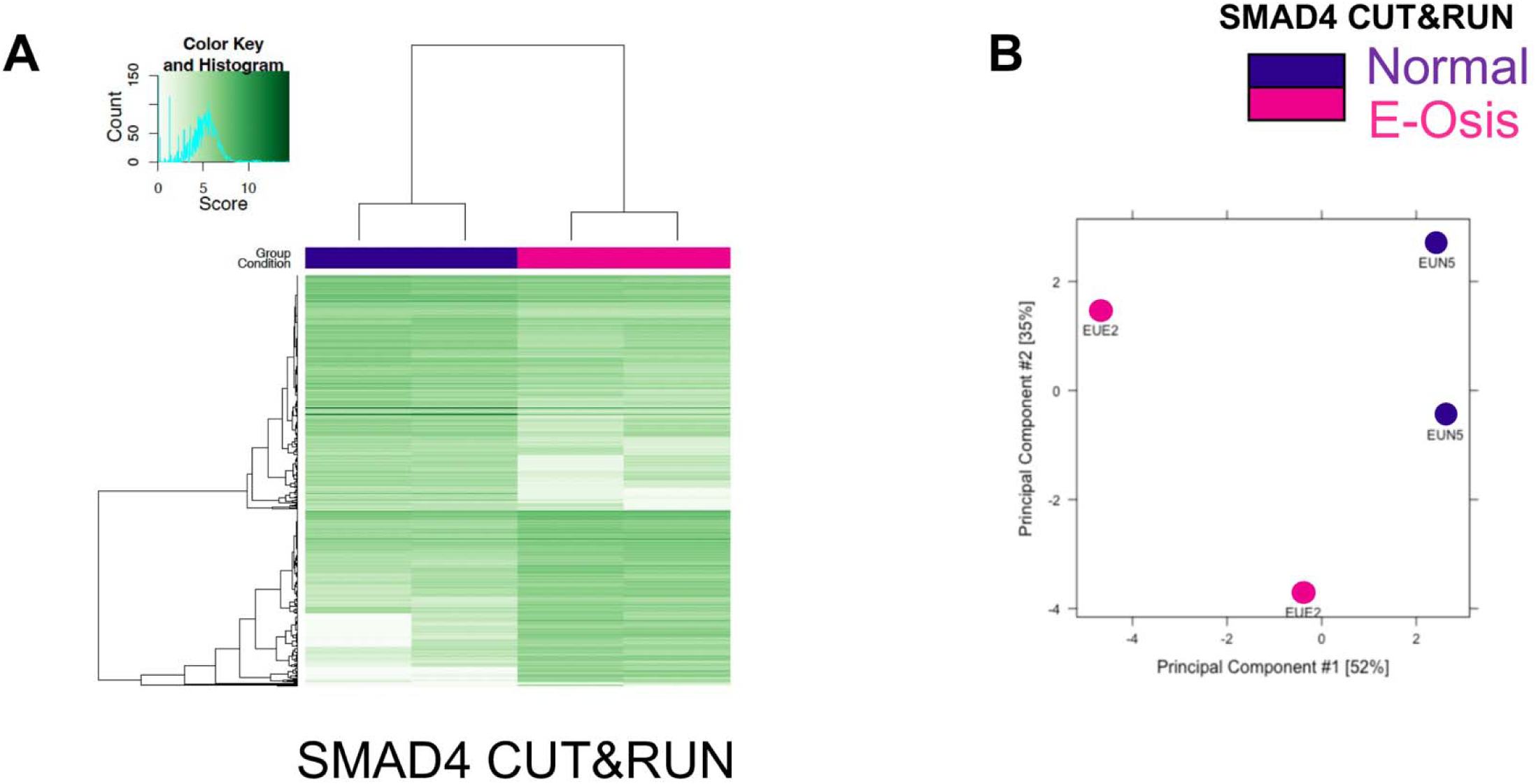
Patient-specific distribution of SMAD4 binding analysis during decidualization. A) SMAD4 CUT&RUN in endometrial stromal cells from individuals with and without endometriosis (E-Osis) after 4 days of EPC treatment. Differential peak signals obtained for the genome-wide SMAD4 distribution in normal versus endometriosis. B) PCA plot of the SMAD4 binding signals comparing the normal versus endometriosis sample replicates.

**Supplemental Figure 4.**
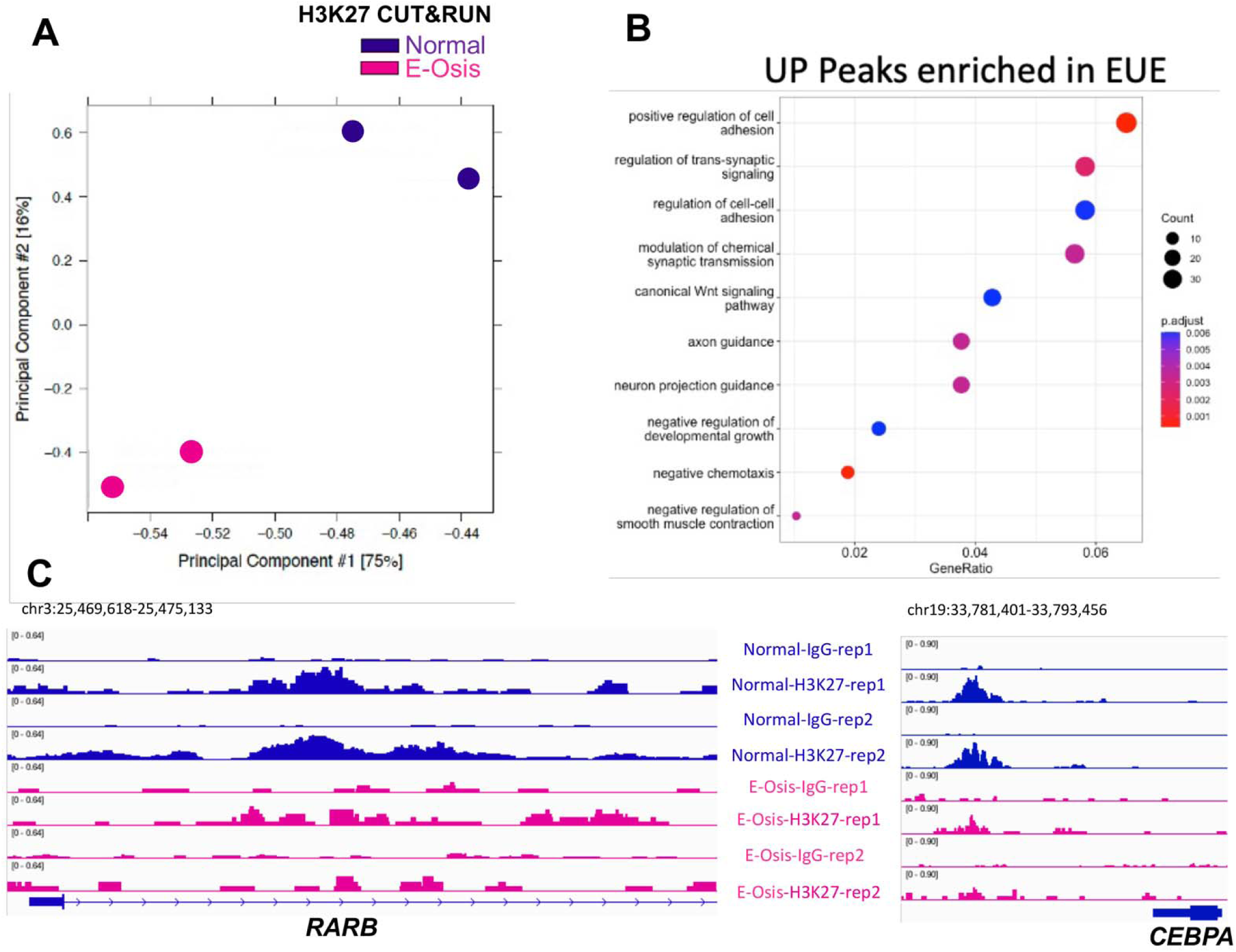
Analysis of H3K27ac in the decidualizing stromal cells of individuals with and without endometriosis. A) PCA plot for H3K27ac replicates of normal and endometriosis stromal cells, showing that the replicates are reproducible. B) Gene ontology classification of the H3K27ac peaks increased in endometriosis stromal cells, showing that categories related to the regulation of cell adhesion were overrepresented. C) Genome track views for the distribution of H3K27ac peaks in the promoter regions of *RARB* and *CEBPA,* showing increased peak density in the endometrial stromal cells from individuals without endometriosis.

**Supplemental Figure 5.**
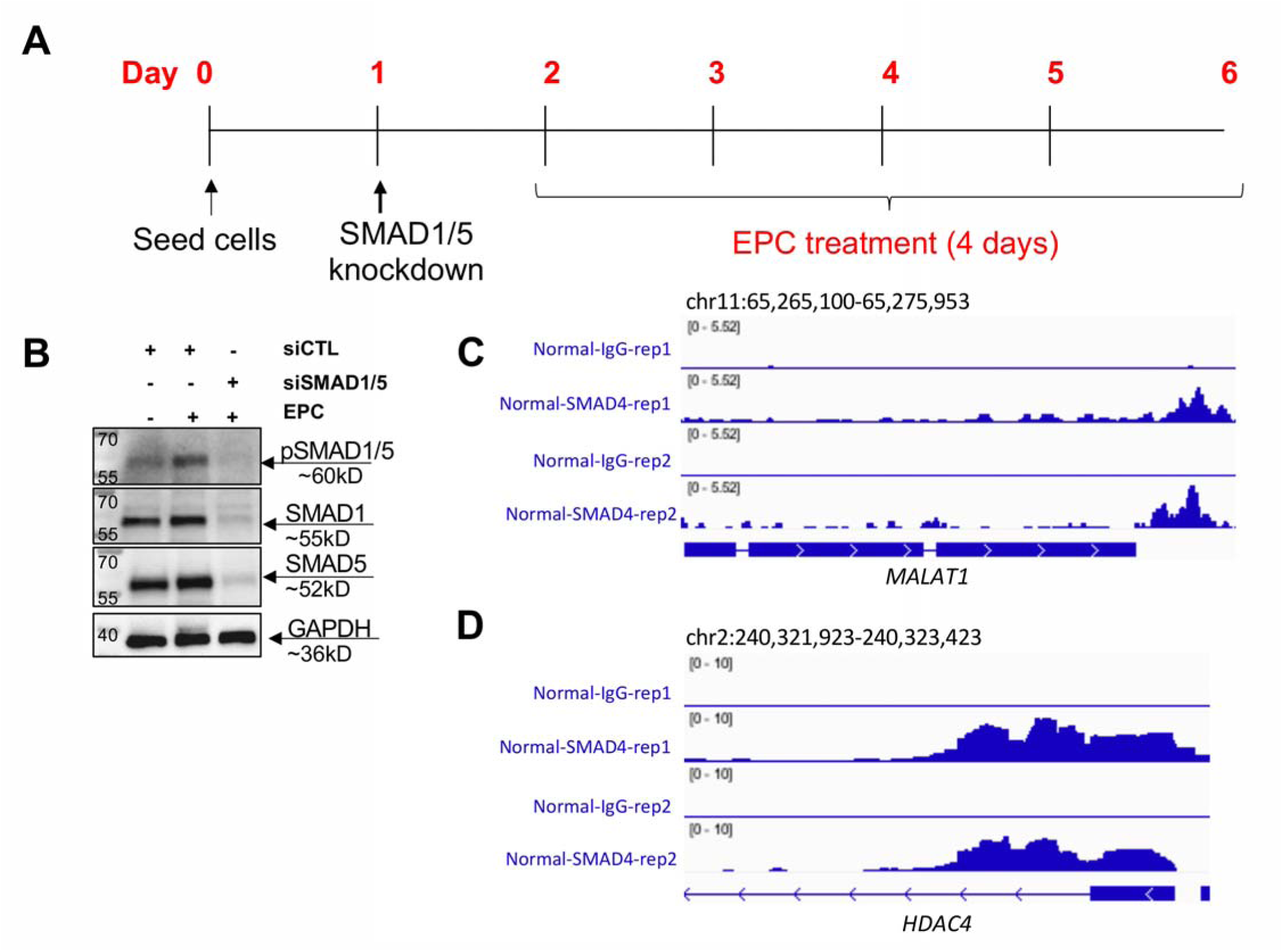
siRNA knockdown of SMAD1 and SMAD5 perturbs decidualization. A) Experimental layout showing the time points at which endometrial stromal cells were plated, transfected with SMAD1/SMAD5 siRNAs, and induced to decidualize with 35nM estradiol, 1µM medroxyprogesterone acetate and 50µM cAMP (EPC) for 4 days. B) Western blot of endometrial stromal cells treated with non-targeting siRNAs (siCTL) or siRNAs targeting SMAD1 and SMAD5 and EPC. Membranes were probed with pSMAD1/5, total SMAD1, total SMAD5 and GAPDH antibodies to confirm knockdown of SMAD1/5 was successful. C-D) Genome track views of the *MALAT1* and *HDAC4* genes showing enrichment of the SMAD4 peaks.

**Supplemental Figure 6.**
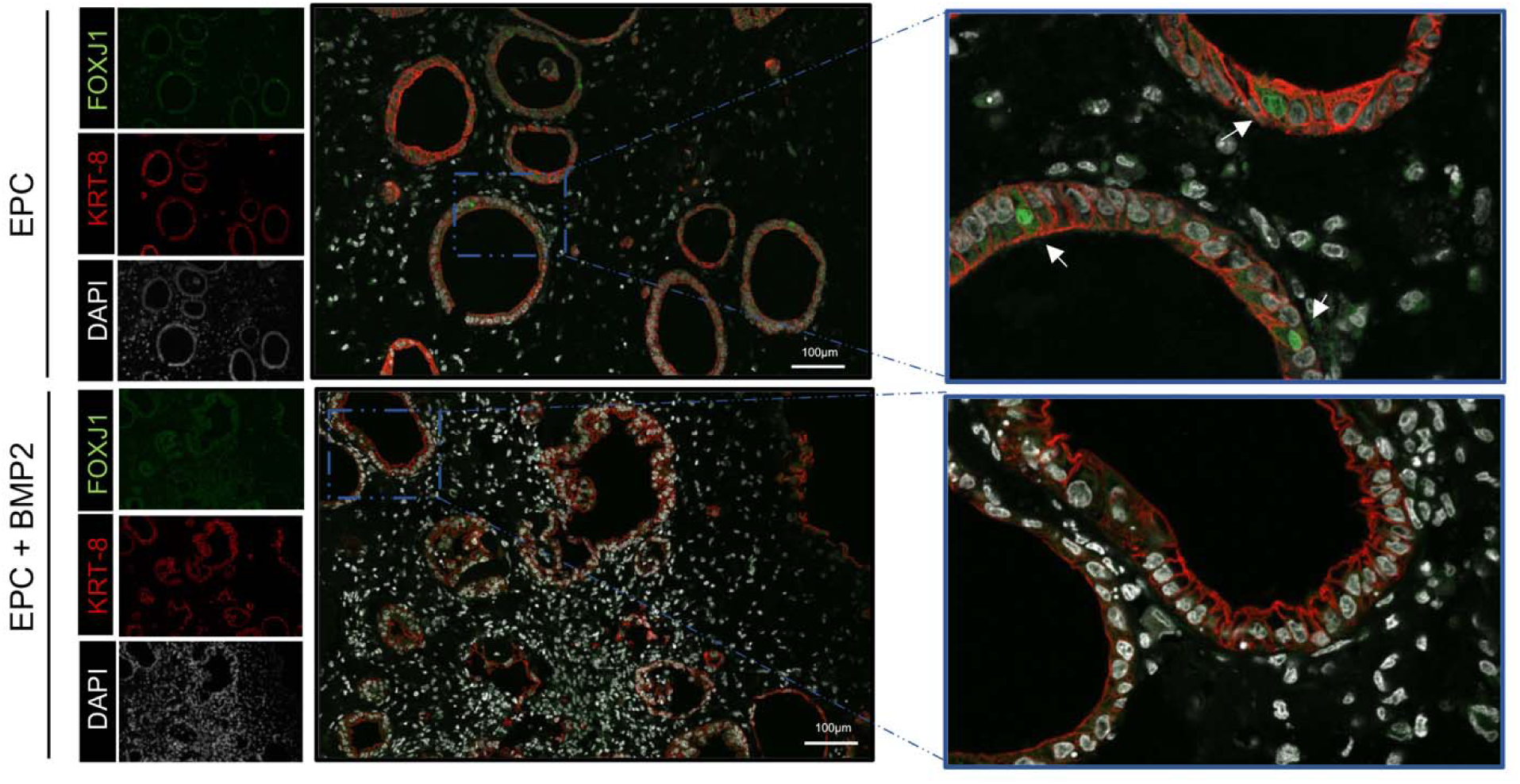
Analysis of the ciliated cell marker, FOXJ1, in decidualizing 3D endometrial assembloids. Immunostaining of the ciliated cell marker, FOXJ1 (green), the epithelial cell marker (cytokeratin 8, KRT-8, red) and DAPI (white) in endometrial assembloids treated with EPC or EPC + BMP2, the hormonal stimuli.

**Supplemental Figure 7.**
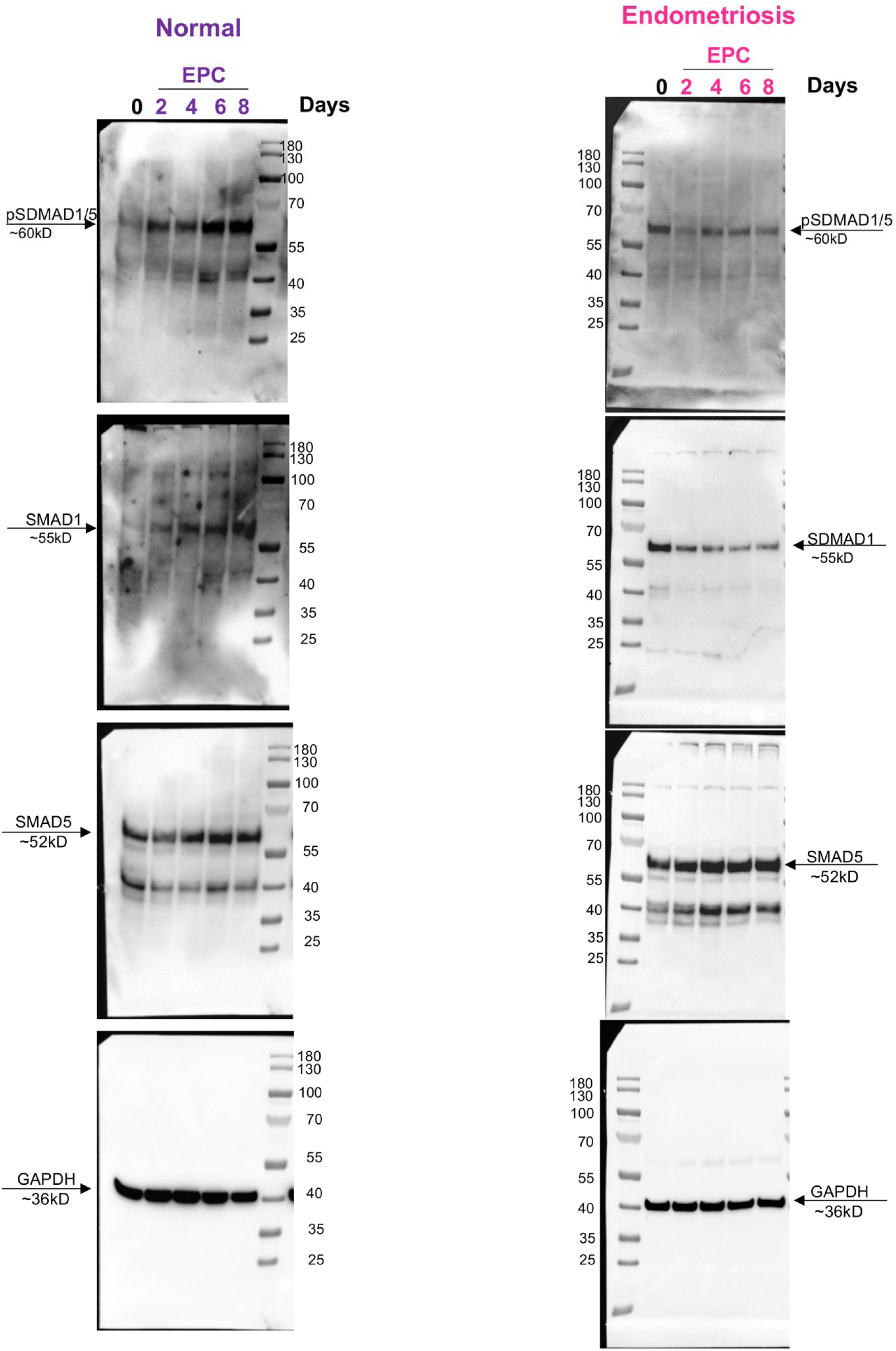
Uncropped western blot images corresponding to Figure 3.

**Supplemental Figure 8.**
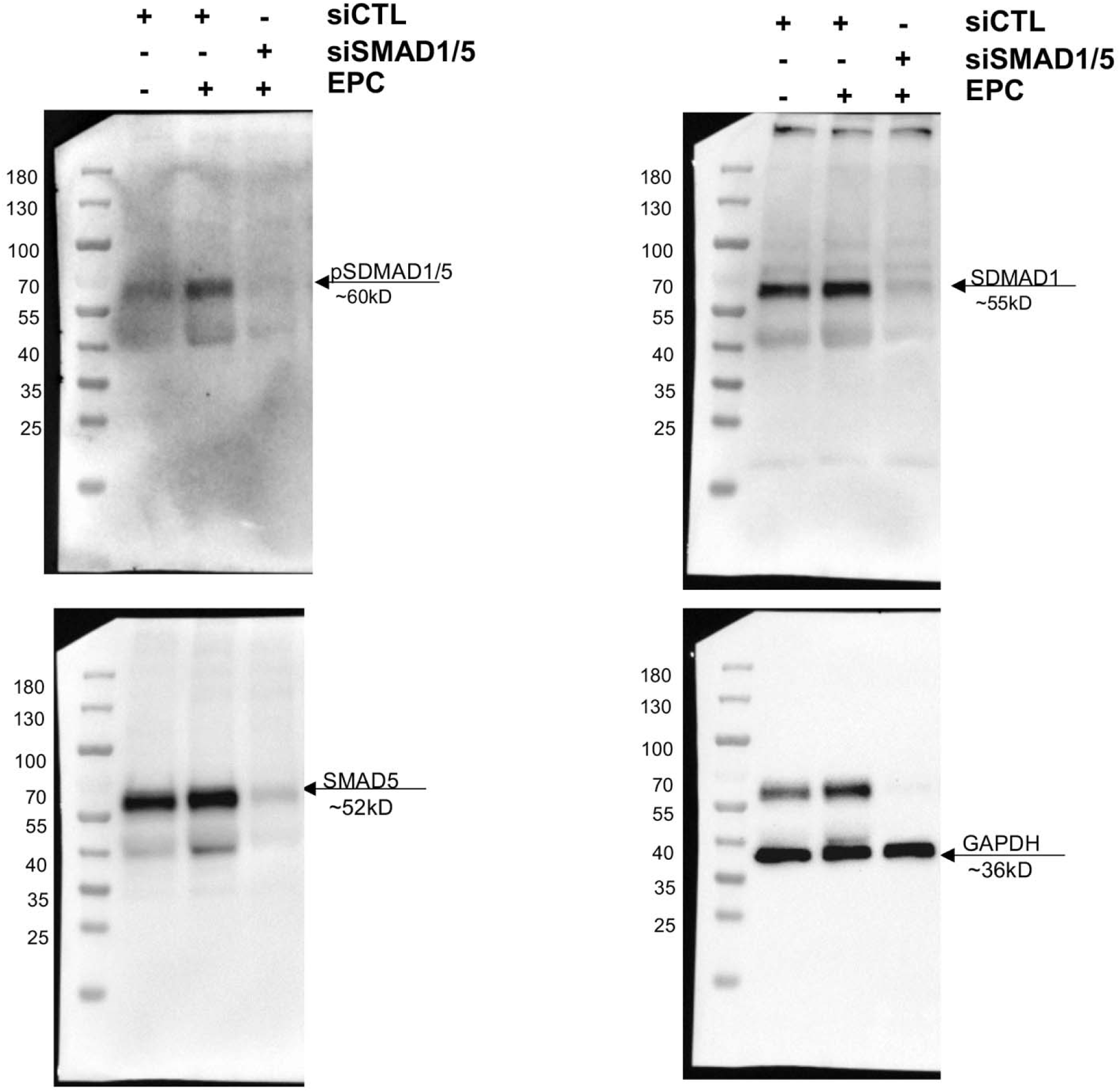
Uncropped western blot images corresponding to Supplemental Figure 5.

**Supplemental Figure 9.**
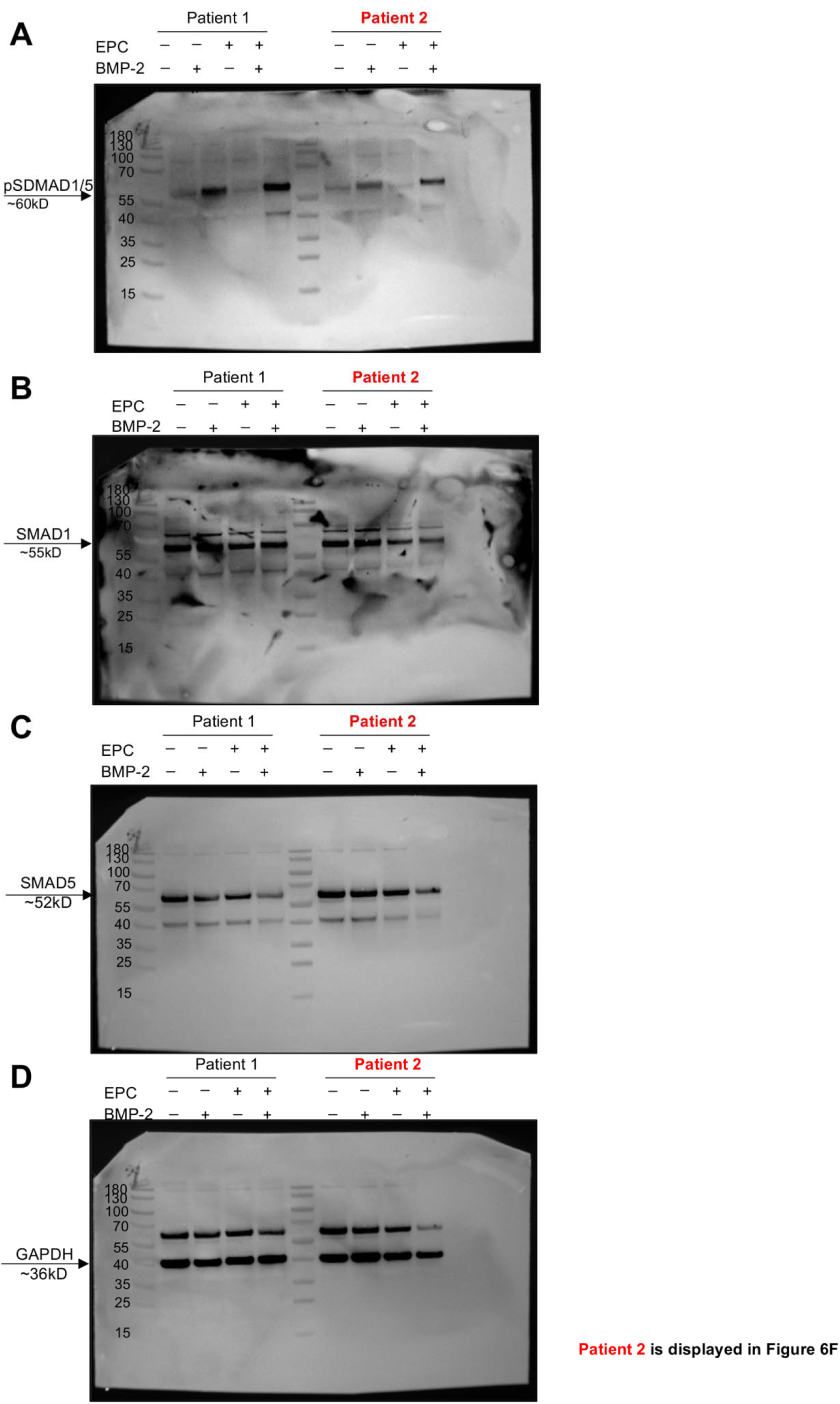
Uncropped western blot images corresponding to Figure 6.

## SUPPLEMENTAL TABLES

**Supplemental Table 1.** Differentially expressed genes in endometrial stromal cells from individuals with and without endometriosis during the time course decidualization treatment compared to baseline Day 0.

**Supplemental Table 2.** Gene ontology classification of differentially expressed genes in the endometrial stromal cells from individuals with and without endometriosis during time course decidualization compared to baseline Day 0.

**Supplemental Table 3.** Transcription factor regulators determined by EnrichR analysis from the genes that were differentially expressed during decidualization in endometrial stromal cells from individuals with and without endometriosis relative to baseline Day 0.

**Supplemental Table 4.** Complete gene expression matrix comparing endometrial stromal cells from individuals with and without endometriosis at each time point during decidualization and the DisGeNET analysis on the 48 consistently down regulated genes.

**Supplemental Table 5.** SMAD4 peak annotations in endometrial stromal cells from individuals with and without endometriosis treated with EPC for 4 days.

**Supplemental Table 6.** Peak annotation of H3K27ac in endometrial stromal cells from with and without endometriosis treated with EPC for 4 days.

**Supplemental Table 7.** Genes that are differentially regulated following EPC treatment and SMAD1/5 siRNA knockdown (siCTL + EPC vs. siSMAD1/5 + EPC).

**Supplemental Table 8.** PCR primer sequences.

**Supplemental Table 9.** List of antibodies and dilutions.

