## Supplemental Table 8 for "Impaired bone morphogenetic protein signaling pathways disrupt decidualization in endometriosis"

**Supplemental Table 8.** qRT-PCR Primer Sequences

| **Gene** | **Forward (5'-3')** | **Reverse (5'-3')** |
| --- | --- | --- |
| *BMP2* | ACCCGCTGTCTCTAGCGT | TTTCAGGCCGAACATGCTGAG |
| *IGFBP1* | TTGGGACGCCATCAGTACCTA | TTGGCTAAACTCTCTACGACTCT |
| *PRL* | AAGCTGTAGAGATTGAGGAGCAAAC | TCAGGATGAACCTGGCTGACTA |
| *FOXO1* | TGATAACTGGAGTACATTTCGCC | CGGTCATAATGGGTGAGAGTCT |
| *WNT4* | CTCCACACTCGACTCCTTGC | CCGAAGAGATGGCGTACACG |
| *SPP1* | TGCAGCCTTCTCAGCCAAA | GGAGGCAAAAGCAAATCACTG |
| *GAPDH* | ACAACTTTGGTATCGTGGAAGG | GCCATCACGCCACAGTTTC |
