## Supplemental Table 9 for "Impaired bone morphogenetic protein signaling pathways disrupt decidualization in endometriosis"

**Supplemental Table 9.** Antibody Lists

| **Antibody** | **Vendor** | **Catalog** | **Application** |
| --- | --- | --- | --- |
| pSMAD1/5 | Cell Signaling | 9516 | WB 1:1000 |
| SMAD1 | Invitrogen | 38-5400 | WB 1:1000 |
| SMAD5 | Proteintech | 12167-1-AP | WB 1:1000 |
| GAPDH | Proteintech | HRP-60004 | WB 1:5000 |
| Vimentin | Cell Signaling | 5741 | IF 1:200 |
| SMAD4 | Abcam | ab40759 | CUT&RUN 1:50 |
| H3K27Ac | Cell Signaling | 8173 | CUT&RUN 1:50 |
| KRT-8 | DSHB | TROMA-I | IF 1:50 |
| FOXJ1 | Sigma | HPA005714 | IF 1:100 |
| Antibody-Rabbit IgG (H+L) Highly Cross-Adsorbed_Alexa Fluor 488 | ThermoFisher | A-21206 | IF 1:250 |
| Antibody-Rat IgG (H+L) Highly Cross-Adsorbed_Alexa Fluor 594 | ThermoFisher | A-21209 | IF 1:250 |
